# Genetic Toolbox for *Photorhabdus* and *Xenorhabdus*: pSEVA based heterologous expression systems and CRISPR/Cpf1 based genome editing for rapid natural product profiling

**DOI:** 10.1101/2024.01.07.574529

**Authors:** Alexander Rill, Lei Zhao, Helge B. Bode

## Abstract

**Background:** Bacteria of the genus *Photorhabdus* and *Xenorhabdus* are motile, Gram-negative bacteria that live in symbiosis with entomopathogenic nematodes. Due to their complex life cycle, they produce a large number of specialized metabolites (natural products) encoded in biosynthetic gene clusters (BGC). Genetic tools for this genus have been rare and applicable to only a few strains. In the past, several tools have been developed for the activation of BGCs and the deletion of individual genes. However, these often have limited efficiency or are time consuming. Among the limitations, it is essential to have versatile expression systems and genome editing tools that could facilitate the practical work.

**Results:** In the present study, we developed several expression vectors and a CRISPR-Cpf1 genome editing vector for genetic manipulations in *Photorhabdus* and *Xenorhabdus* using SEVA plasmids. The SEVA collection is based on modular vectors that allow exchangeability of different elements (e.g. origin of replication and antibiotic selection markers with the ability to insert desired sequences for different end applications. Initially, we tested different SEVA vectors containing the broad host range origins and three different resistance genes for kanamycin, gentamycin and chloramphenicol, respectively. We demonstrated that these vectors are replicative not only in well-known representatives, e.g. *Photorhabdus laumondii* TTO1, but also in other rarely described strains like *Xenorhabdus* sp. TS4. For our CRISPR/Cpf1-based system, we used the pSEVA231 backbone to delete not only small genes but also large parts of BGCs. Furthermore, we were able to activate and refactor BGCs to obtain high production titers of high value compounds such as safracin B, a semisynthetic precursor for the anti-cancer drug ET-743.

**Conclusions:** The results of this study provide new inducible expression vectors and a CRISPR/CPf1 encoding vector all based on the SEVA (Standard European Vector Architecture) collection, which can improve genetic manipulation and genome editing processes in *Photorhabdus* and *Xenorhabdus*.

## Background

Microorganisms often use low-molecular weight compounds to interact with other organisms in complex environments [1-3]. These compounds, also known as secondary or specialized metabolites or natural products can be structurally and functionally diverse [4]. Their ecological function ranges from defense against competitors or enemies [5], cell-cell communication, to symbiosis and pathogenesis [6, 7]. These functions also allow their application in medicine, for example, as anticancer or antibacterial drugs [8]. Increasing numbers of infectious diseases caused by bacteria and viruses make it necessary to constantly find new natural products serving as drug candidates also against resistant pathogens. Since climate change as well as other human impact on our planet also plays an important role in the occurrence of such infectious diseases, the demand for such compounds likely will even increase in the future [9, 10].

The progress in sequencing technology opens up new possibilities to study the genomic and metabolic diversity of microorganisms [11]. In addition, there are numerous tools and databases for the prediction of genes related to the biosynthesis of secondary metabolites that enable simple and accurate genome mining [12, 13]. In general, the predicted number of biosynthetic gene clusters (BGCs) responsible for the production of natural products in any genome outcompetes the number of produced natural product types observed under laboratory conditions [14]. Therefore, robust and versatile molecular tools need to be developed to activate BGCs that might be silent under laboratory conditions. One possibility is to express the BGC heterologous in related organisms [15]. For this purpose, several methods have been established to assemble large DNA fragments, such as TAR cloning (Transformation-Associated Recombination) and ExoCET, which is based on the Red/ET recombineering tool [16-19]. Another way to activate BGCs is to manipulate the regulatory network in the native host. Since promoters are the first target for regulatory mechanisms, methods have been developed to increase their strength, replace them with a set of regulatory sequences with known functions resulting in the exchange of the natural against an inducible promoter [20] or to prevent binding of repressors to the promoter sequence [21]. Various methods have been developed for this purpose, such as a phage recombinase-based knock-in approach, in which an antibiotic resistance gene including promoter was integrated in front of a gene cluster [22, 23]. The increasingly common method of CRISPR/Cas also offers a good opportunity for seamless and precise chromosomal integration or replacement of heterologous sequences for BGC activation [24].

Bacteria of the genus *Photorhabdus* and *Xenorhabdus* are a prolific source for several natural products [4, 25, 26]. Due to their particular life cycle that involved interaction with a nematode host, insect prey and other microbes and organisms as competitors, they produce natural products showing activity in cell-cell communication or antibiotic, antifungal, insecticidal and cytotoxic activity [3, 27-31]

Molecular tools have also been developed for *Photorhabdus* and *Xenorhabdus* to elucidate new natural products and genes involved. The easy Promoter Activated Compound Identification (easyPACId) method is based on the integration of a non-replicating plasmid containing an R6K ori with a heterologous promoter in front of a BGC. Coupled with the deletion of the gene encoding the global regulator Hfq, the production of individual compound classes from one BGC in an almost clear metabolic background can be achieved [32]. A CRISPR/Cas9 system was also established recently for *Photorhabdus laumondii* TTO1. Based on Cre-lox recombinase-mediated genome engineering and stepwise integration of the Cas9 gene and sgRNA into the chromosome of the target organism, subsequent integration of individual sgRNAs and repair donor templates allows for sequential deletion of genes of interest [33, 34].

In addition to the well-established type II CRISPR/Cas9 system, the use of a type V-A CRISPR-Cas system (CRISPR-Cpf1) is becoming increasingly widespread. Unlike Cas9, only a single crRNA is required and the *cas12* gene is smaller than *cas9*, while it recognizes a T-rich PAM sequence [35].

Here we report the development of pSEVA based expression systems with a variety of replication origins, resistant genes and promoters. Furthermore, we developed an efficient and simple genome editing strategy using a CRISPR/Cpf1 system assisted by λ red recombination encoded on a single “all-in-one” plasmid (pAR20) in *Photorhabdus laumondii* TTO1, *Xenorhabdus nematophila* and *Xenorhabdus* sp. TS4. Besides deletions and promoter exchanges, we apply our approach to the activation and optimization of the previously described and characterized safracin B (SAC-B, **16**) in *Xenorhabdus sp*. TS4 up to a final production titer of 336 mg/L. Safracins have been previously described and characterized in *Pseudomonas fluorescens* [36, 37]. They belongs to the tetrahydroisoquinoline (THIQ) family natural products with unusual scaffolds and bioactivities [38]. The most prominent of the THIQ family, ET-743, has already been approved by the Food and Drug Administration for the treatment of soft tissue sarcomas [39]. However, ET-743 is difficult to obtain directly from its natural source [40]. Therefore, extensive synthesis studies have been performed [41] although the main industrial production is by semi-synthesis using cyanated safracin B as starting material [42, 43].

## Methods

### Strains, plasmids, and growth conditions

All bacterial strains and plasmids used in this study are listed in Table S1. *E*. *coli* DH10B were used as host for plasmid construction, *E*. *coli* ST18 was used as donor strain for conjugation to transfer plasmids into *Photorhabdus laumondii* subsp. *laumondii* TTO1 and *Xenorhabdus nematophila* ATCC 19061. *E. coli* was cultured in LB medium at 37 °C, and *Photorhabdus/Xenorhabdus* was cultured in LB medium at 30 °C. All liquid cultures were incubated at 200 rpm and antibiotics were added as follows: 50 µg/ml kanamycin (Km), 50 µg/ml chloramphenicol (Cm) and/or 50 µg/ml gentamycin (Gm) for *E. coli*. 25 µg/ml kanamycin, 15 µg/ml chloramphenicol and 7.5 µg/ml gentamycin were used for *Photorhabdus/Xenorhabdus*. 1.5 g/L agar was added to LB for agar plates. In addition, the concentration of L-arabinose (Ara) and anhydrotetracycline (AHT) to induce expression of the Cpf1 protein and lambda red proteins was 0.4% (w/v) and 200 nM, respectively. For inducing of genome-integrated promoters, 100 µM of vanillic acid (VA) and 1 mM IPTG was used. Plasmid based expression was induced with 0.2% (w/v) L-arabinose, 100 µM vanillic acid, 1mM IPTG and 100 µM cumaric acid, respectively. All strains were kept as glycerol stocks prepared in LB containing 25% glycerol at −80 °C.

### Construction of editing plasmids

All plasmids used in the study are listed in Table 1 in Additional File 2. Sequences of the crRNAs, primers and synthetic DNA fragments used in the study are in listed in Additional File 2. pAR16 harboring *cpf1* and lambda red genes harboring vector was constructed by exchanging of the origin of replication via NEB Hifi assembly. AR426/427 was used for backbone amplification from p46Cpf1-OP2 (obtained from Addgene) and AR428/429 was used to amplify the p15A ori from a pACYC-derived vector. pAR18 was constructed in three steps: 1) pTargetF (obtained from Addgene) was linearized with the primer pair AR430/431 while the *cas9* sgRNA sequence was replaced by the *sacB* gene including its native promoter (amplified with AR432/433 from pEP17-KM [44]); the final pAR14 was assembled via NEB Hifi assembly. 2) To exchange the spectinomycin resistance, pAR14 was linearized by restriction digestion with *Xho*I and *Sac*I; the desired fragment was purified via gel extraction (Monarch® DNA Gel Extraction Kit) and assembled with the gentamycin resistance gene amplified from pSEVA631 (AR434/435) to obtain pAR15; 3) In the last step, the Cpf1 crRNA framework encoded on a synthetic gene fragment (synthesized by GENEWIZ Germany GmbH) was inserted into the linearized pAR15 (AR436/437) via NEB Hifi assembly, giving pAR18.

**Table 1.**
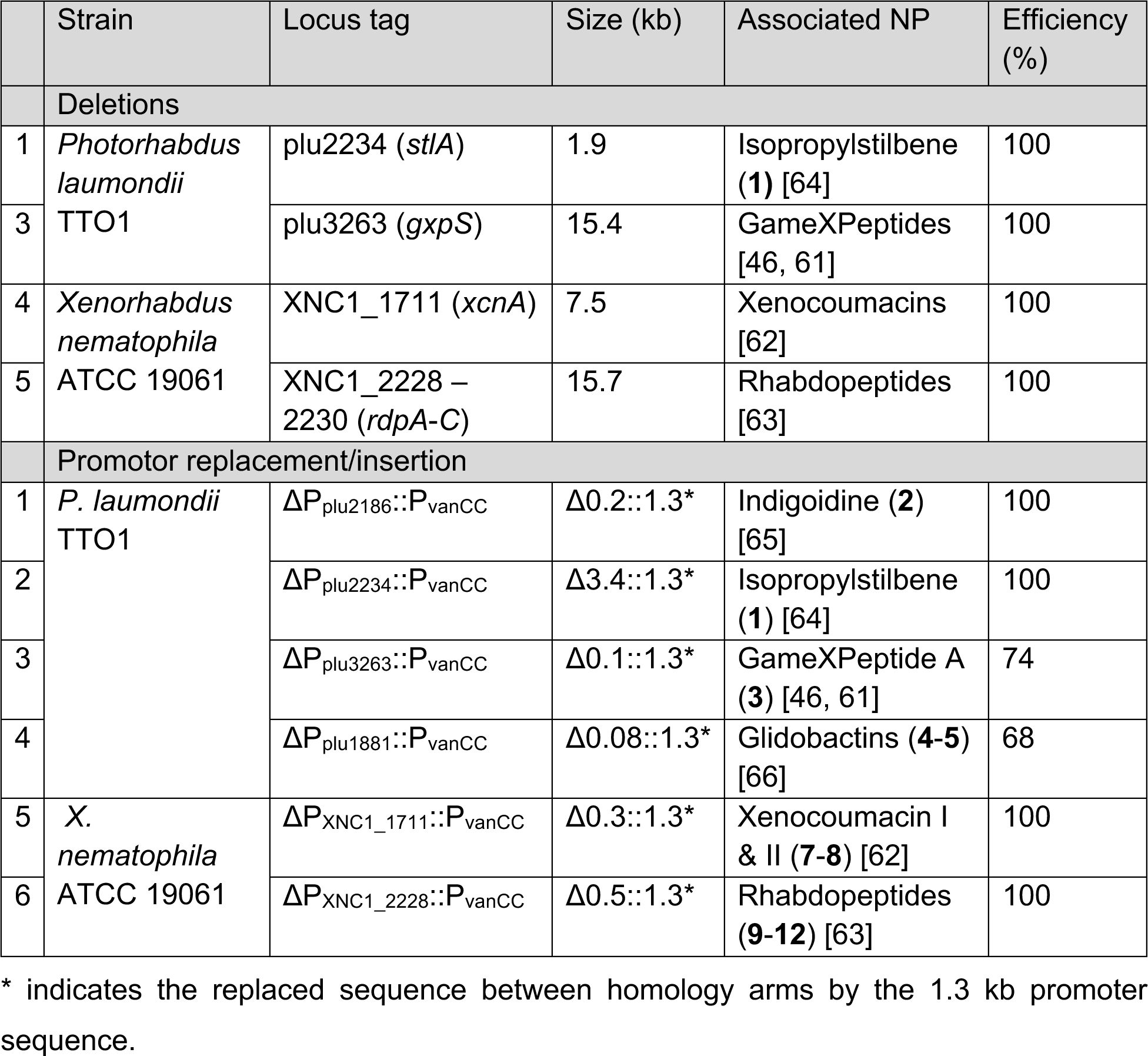
Overview of successful gene/ BGC editing.

The empty pAR20 was constructed *in silico* and synthesized as whole plasmid by Double Helix Technologies (DOULIX™). pAR20 editing plasmids were assembled via NEB Golden Gate Assembly. Target sequence and homology repair templates, ordered as synthetic dsDNA gene fragments, and were obtained from Twist Bioscience and Integrated DNA Technologies, respectively. The NEB Golden Gate Assembly mix containing pAR20 and dsDNA inserts was prepared according to the manufacturer’s instructions. Correct assembly was checked by colony PCR using primer pair AR533/534.

### Electrotransformation and biparental conjugation

Electrocomponent *Photorhabdus* and *Xenorhabdus* cells were prepared as previously described [17]. For conjugation of the pAR20/pSEVA derivatives overnight cultures of *E*. *coli* ST18 and *Photorhabdus*/*Xenorhabdus* were used, respectively. The optical density of the two cultures was measured and set to a ratio of 4:1 Recipient/Donor in a total volume of 1 ml. The cell mixture was centrifuged at 5000 x g and washed twice with 1 ml LB. Finally, the cell pellet was resuspended in 100 µl LB and spotted onto a LB agar plate containing 50 µg/ml aminolevulinic acid. The next day, the cell mass was scraped off and resuspended in 1 ml LB. A dilution of 1:50 was plated on a selection plate with appropriate antibiotics.

### Cpf1 assisted gene editing with pAR20 series and plasmid curing

For general gene editing, a strain carrying a gene-specific pAR20 vector was inoculated as a 5 ml overnight culture in a 50 ml Erlenmeyer flask. The next day, this culture was diluted to a final OD of 0.5 in 10 ml LB containing 25 µg/ml kanamycin. The culture was then incubated at 30 °C until an OD of 0.8 - 1.0 was reached. After addition of AHT, the culture was further incubated at 25 °C for 1h.

Subsequently, L-arabinose was added and incubated for another 3 h at the same temperature. Finally, 50 µl of the cell suspension was plated on a selection plate with kanamycin and arabinose. Successful editing approaches were confirmed by colony PCR [45].

To generate a plasmid free stain a 1:500 dilution of the overnight culture was inoculated into fresh LB culture containing 10% sucrose and incubated for further 20 h. This culture could be used directly for further rounds of genome editing.

### Construction of expression plasmids

To generate pSEVA-based expression vectors, the empty pSEVA plasmids were amplified with the primer pair AR1081/AR900. The resulting PCR fragments were assembled with NEBuilder® HiFi DNA Assembly Master Mix together with mNeonGreen synthetic dsDNA fragment and synthetic dsDNA fragments of the selected promoters. For further use as an expression vector template, the cloned plasmids were amplified with the primer pair AR900/970 and assembled with the desired insert fragment.

### Construction of plasmids for expression of safracin derivatives

For construction of the expression plasmids in *E. coli mtaA* pCOLA, pACYC and pCDF empty backbones were linearized by restriction digest with *NcoI* and *PstI*. Inserts were amplified by PCR with the according primer pairs (Additional File Table 1) and purified via gel extraction (Monarch® DNA Gel Extraction Kit). Purified PCR fragments were assembled with the corresponding plasmid backbones by HiFi cloning (NEB; E2621L). Subsequently the assembly mix was transformed by electro transformation into *E. coli mtaA*. Positive clones were verified by restriction digest.

### Safracin A and B production

For the production of safracin A and B a single colony of the selected strain was inoculated in 5 ml LB with appropriate antibiotics for overnight cultivation. After 18h, the overnight culture was diluted 1:50 in 45 ml fresh LB medium with antibiotics and incubated at 30 °C until OD 0.8 was reached. Subsequently the culture was centrifuged in a 50 ml centrifugation tube at 20 °C and 4000 rpm. The supernatant was discarded and the cell pellet was resuspended in 15 ml fresh modified Bacto™ CD Supreme Fermentation Production Medium (supplemented with 1 mM L-tyrosine, 1 mM L-methionine, 0.5 g/L L-leucine, 1 g/L pyruvate, 1 ml/L trace element solution as described in [32], 2 ml/L vitamine solution as described in [32] and 1% (w/v) sodium succinate as carbon source). Inducers for BGC activation and Amberlite™ XAD16N resin (2% final) were added after 1 h cultivation at 28 °C. The Amberlite™ XAD16N resin was harvested after 3 day of cultivation and extracted with a culture volume equivalent of methanol. Methanol extracts were centrifuged at 13,000 rpm for 30 min and transferred to HPLC vials at a final dilution of 1:100. UPLC-HRMS analysis of extracts, MS-MS fragmentation analysis and labelling experiments for MS-based structure elucidation of new natural products were performed as described previously [46, 47]. SAC-B concentrations were quantified by serial dilution and resulting standard curve of a standard of SAC-B obtained from Focus Biomolecules (CAS: 87578-99-2, Catalog Number: 10-3208). SAC-A was also quantified using this standard curve.

### Production and isolation of safracin derivative 20

The compound **20** was isolated from heterologous expression of the safracin homologous gene cluster containing the genes xets_03193-03202 without the peptidase-encoding gene xets_03192 from Xenorhabdus TS4 in *E. coli mtaA*. The production of compound **20** by the heterologous *E. coli* was confirmed by HPLC-MS analysis. To isolate compound **20**, overnight cultures of the heterologous *E. coli* were inoculated into LB medium with appropriate antibiotics and 0.1% L-arabinose (from a 25% stock solution) added. The cultures were cultivated at 30 °C and 200 rpm on a rotary shaker. The cells were harvested after 3 days and extracted with MeOH by ultrasonic method for 15 min. The extract was purified by semipreparative HPLC system using a gradient MeCN/H_2_O solution containing 0.1% formic acid to yield the compound **20**. The structure of the compound **20** was confirmed by HR-MS m/z 763.4994 [M + H]^+^ (calc. for C_44_H_67_N_4_O_7_, 763.5004; Δppm 1.35) and NMR (Table S4 and Figures S8-12). NMR spectra for purified compounds were measured in DMSO-*d_6_*.

### Isolation and Analysis of other natural products

For production of all other natural products, *Photorhabdus/Xenorhabdus* strains were cultured in XPP medium [32] at 28 °C and 200 rpm for 3 days. Subsequently the culture was extensively mixed 1:1 with methanol for 5 min. The HPLC/MS samples were prepared as described above.

## Results and discussion

### Construction of conjugatable expression vectors for *Photorhabdus* and *Xenorhabdus*

To create a conjugatable library of expression vectors for *Photorhabdus* and *Xenorhabdus*, we used pSEVA plasmids obtained from seva-plasmids.com as a template. Their architecture allows to quickly adapting the origin of replication and resistance gene to the desired strain. We linearized the vectors and assembled them with modified promoters (*vanR*/P_vanCC_, l*acI*/P_tac_, *cymR*/P_cymRC_ and *araC*/P_Bad_) as described previously [48]. To test their functionality, we additionally assembled the gene encoding the fluorescence protein mNeonGreen on our expression vectors [49]. The vectors were introduced into the target strains by conjugation and tested for expression of mNeonGreen by addition of the inducers directly into the LB-Agar. We therefore were able to generate a library of 26 vectors with the resistance genes for gentamycin, kanamycin and chloramphenicol as well as the replication origins pBBR1, p15A, pUC, RK2, RSF1010 and pRO1600/ColE1 (Table S1, Plasmids).

### Optimization of the two-plasmid-based CRISPR/Cpf1 method for *Photorhabdus*

The CRISPR/Cpf1 editing method described here is based on the method described previously [50]: A helper and a donor plasmid are used to perform genome editing supported by the lambda recombination system. The helper plasmid p46Cpf1-OP2 encodes an *E*. *coli* codon-optimized variant of the *cpf1* gene under the P*_Bad_* promoter (inducible with L-arabinose (Ara)) and the lambda red (*gam*, *exo*, *bet*) recombination system controlled by a P_tet_ promoter (inducible with anhydrotetracycline (AHT)). The donor plasmid is based on a high copy pUC vector carrying the crRNA sequence for a specific target gene, as well as homology regions to the target sequence serving as repair templates after double strand break.

In order to validate this method in *Photorhabdus laumondii* TTO1, the *stlA* gene (*plu2234*) was selected as the first target for genome editing because of an easily recognizable phenotype (Fig. S1 A). StlA is a phenylalanine ammonia lyase involved in the biosynthesis of isopropylstilbene (IPS; **1**) produced by all *Photorhabdus* strains (Fig. S1 A) and its deletion results in overproduction of the orange anthraquinone pigments [51].

We constructed a new donor plasmid (pAR18, Fig. S1 B) based on the pTargetF vector with gentamycin resistance as well as the *sacB* gene as counter-selection marker enabling rapid loss of the donor plasmid allowing for several rounds of genome editing. Unlike the original method, we decided to use two spacer sequences for each deletion to enable targeting of the leading and lagging strand, respectively. For this purpose, we constructed a synthetic framework consisting of the constitutive promoter J23119, a transcriptional terminator and the direct repeats of the crRNA sequence. We modified the described CRISPR array [35] and replaced the spacer sequence between the FnCpf1 specific direct repeats by two *Bsa*I restriction sites containing sequence (Fig. S1B). The *stlA* specific spacer sequences were determined by “Annotate & Predict” function of Geneious Prime, “TTTN” was set as PAM site and a spacer sequence length of 31 bp. Furthermore, to facilitate plasmid assembly, the homology repair arms (HR-L/HR-R; ∼500 bp) were each coupled with one of the crRNAs and synthesized as a dsDNA fragment (Fig. S1 B; dsDNA/ dsDNA-R). The resulting *stlA*-specific pAR18, assembled by the Golden Gate reaction, was transformed into *Photorhabdus laumondii* TTO1 by electroporation (Fig. S1 B). However, we were not able to transform the original p46Cpf1-OP2 [50] plasmid into *Photorhabdus.* Although it was previously described in [52] that the pSC101 origin of replication replicates in *Photorhabdus*, we assume that the low copy [53] number combined with the chloramphenicol resistance is not suitable for the cells. Therefore, we switched the origin of replication from pSC101 to p15A, which is distinguished by a higher number of copies per cell [53], resulting in the successfully transformable pAR16 (Fig. S1B).

To determine the optimal starting OD_600_ of the liquid CRISPR culture we selected OD = 0.1, 0.2 and 0.5 after inoculation from an overnight culture. Since *Photorhabdus* has a much slower doubling time compared to *E*. *coli*, the required OD_600_ of 0.8-1.0 could not be achieved within a single working day when starting OD_600_ was below 0.5. Therefore, 0.5 was chosen as the starting OD_600_ of the liquid culture for the following experiments. When the OD_600_ of 0.8-1.0 was reached, the inducers for *cpf1* (Ara) and lambda red (AHT) were added. As the system was previously tested [17, 50], we did not check the individual components of the system but only the overall method. To reduce cell growth and promote protein expression [54-56], we set the temperature to 25°C after induction and observed an editing efficiency of 100%, the efficiency decreases at higher temperatures (Fig. S1 C).

### Construction and optimizing of the single-plasmid approach

Since we observed that *Photorhabdus* had poor transformation efficiency and *Xenorhabdus* was not transformable at all with the p46Cpf1-OP2 and its derivative pAR16, we decided to develop a single plasmid solution that had previously been described as particularly efficient in other strains [57-59]. Based on our recently gained knowledge about the highly mobile pSEVA plasmid family we selected the broad-host vector pSEVA231 [60]. We constructed a plasmid (pAR20) combining the components from the donor plasmid pAR18 and helper plasmid p46Cpf1-OP2. A major advantage of the method is the possibility to conjugate the plasmid directly from *E*. *coli* into the recipient strain without any integration of the plasmid or its parts into the genome.

To test its efficiency, we again deleted *stlA* and found all colonies edited and showing the expected pigmentation phenotype. In addition to deleting smaller genes, we also wanted to evaluate the potential of the method for medium and large deletions and whether we could speed up the editing process. We focused on genes whose manipulation have visible phenotypes or result in metabolite changes easily detectable by mass spectrometry. The selected and successfully deleted genes/genome segments include *stlA* (1.9 kb deletion), *gxpS* (15.4 kb deletion) in TT01 and *xcnA* (7.5 kb deletion) and *rdpABC* (15.7 kb deletion) in *Xenorhabdus nematophila* involved in the biosynthesis of GameXPeptides [46, 61], xenocoumacin [62] and rhabdopeptides [63], respectively (Table 1, Figure S3).

Efficiency was determined by dividing the number of positive edited colonies by the total number of colonies tested, pictures of agarose gel electrophoresis are shown in Figure S5. Target sequences used for each approach are listed in Table S2. Large gene deletions up to 15 kb in *P. laumondii* TTO1 and *X*. *nematophila* were easily obtained with high efficiency. Furthermore, we found no major impact on the metabolic profile after deletion of *gxpS* in *P. laumondii* TTO1 or *xcnA* in *X*. *nematophila* except the expected loss of GameXPeptides and xenocoumacins, respectively. Deletion of the *rdpABC* genes responsible for the production of rhabdopeptides (**9**-**12**) in *X*. *nematophila* however also leads to an additional loss of xenortide A (**6**) as well as an increase of xenocoumacin II (**8**) (Fig. S3 B). These findings indicate a regulatory or metabolic network of natural products in *X. nematophila* associated with rhabdopeptide biosynthesis that will be studied in the future.

### Metabolite profiling by insertion/replacement of regulatory sequences

In addition to deletions, the use of small regulatory sequences like inducible promoters is particularly important in natural product research and for biosynthesis elucidation. For this purpose we modified and tested various inducible promoters described in [48] featuring an insulator (*riboJ*) and a strong ribosomal binding site (RBS) to switch on and off different genes and gene clusters. As example, we selected *indC*, encoding a NRPS involved in the biosynthesis of the blue pigment indigoidine [65]. This gene is silent under laboratory conditions but can be activated by the introduction of a foreign promoter [65]. For our approach we “selected” 500 bp upstream of the *indC* coding sequence (HR-L) and 500 bp downstream of the start codon (HR-R), while the distance between both can be variable (Fig. 1A). In contrast to a deletion, a promoter (P_vanCC_) sequence was assembled between the homology arms during Golden Gate Assembly (Fig. 1 A). After the editing approach, the sequence between the homology arms was replaced by the promoter sequence.

**Figure 1.**
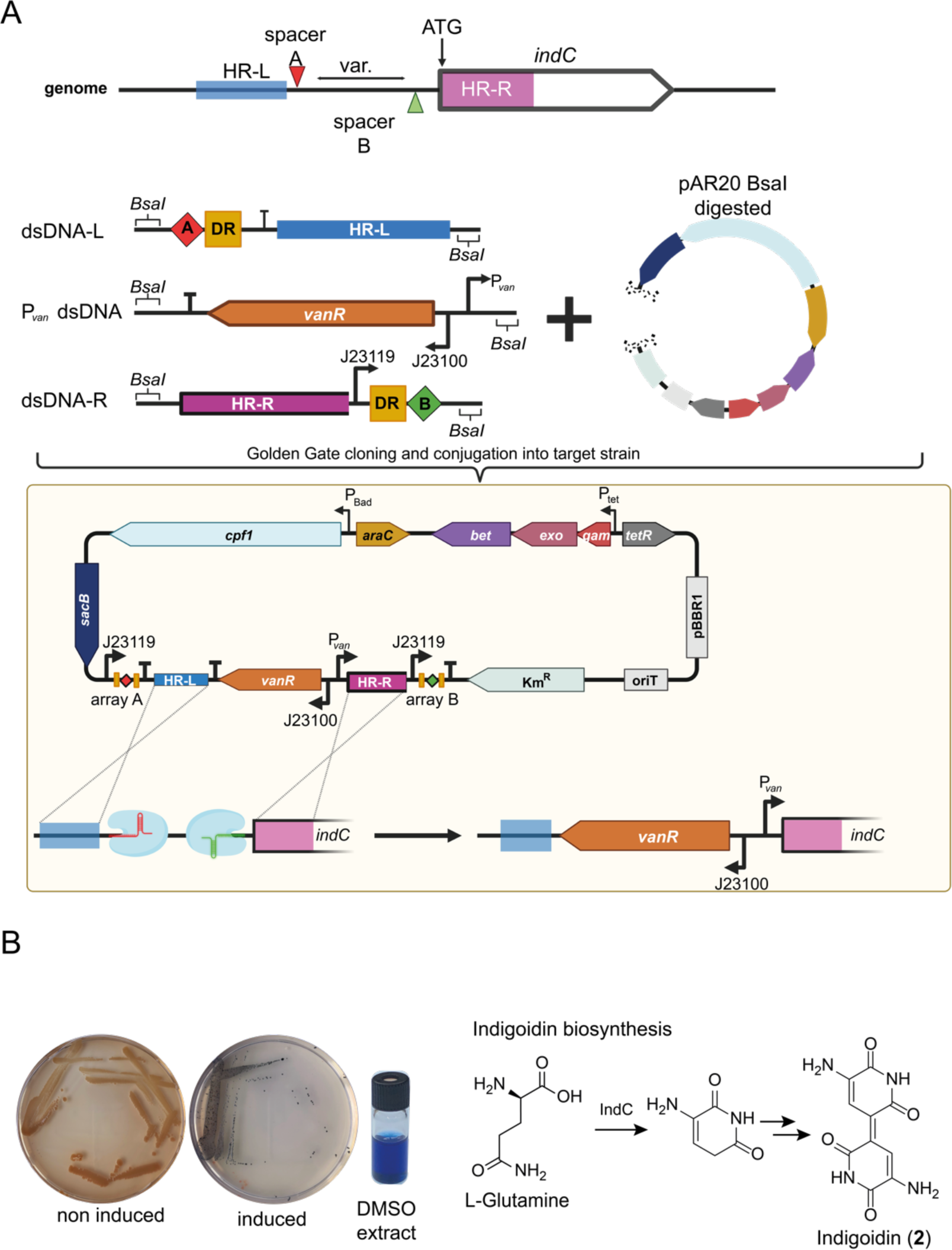
Single plasmid CRISPR/Cpf1 genome editing method for promoter exchange upstream of *indC*. **A**) Schematic representation of spacer sequence and homology regions selection; schematic cloning of synthetic dsDNA fragments harboring the repair templates and spacer sequence and the vanillic acid inducible promoter sequence (P*_van_*) into BsaI digested pAR20; schematic representation of the genome editing approach mediated by pAR20; the sequence between the homology arms is replaced by the promoter sequence; detailed genotype of pAR20 can be found in Table S1. HR-L/R = homologues region L (left) or R (right), dsDNA = synthetic double strand DNA, DR = FnCpf1 specific direct repeat, red or green diamond = spacer sequence A or B, crossing dashed lines representing crossover event after CRISPR induced double strand break. **B**) Phenotypic comparison of induced and non-induced mutants; DMSO extract of induced mutant; biosynthesis of **2.** Figure was taken from BioRender.com and modified.

We tested the colonies by PCR and achieved 100% editing (Fig. S5). After induction of the promoter with vanillic acid, the typical blue pigment was observed (Fig. 1 B). We also applied this approach to other genes of known BGCs facilitating overexpression of previously moderate produced metabolites as well as production of previously “silent” BGCs. The successfully edited targets are listed in Table 1 including six different compound families. MS spectra of induced and non-induced cultures after promoter insertion are shown in Figure S4. In general, a significantly higher production of natural products was observed through compared to the wild type level. As previously observed during *rdpABC* deletion, the promoter exchange upstream of *rdpA* had an impact on xenortide A (**6**) production. While **6** was still detectable in the non-induced culture, it was significantly reduced after induction of the *rdp* BGC, which might be due to the cross talk between RdpABC and XrdAB as shown in Fig. S3.

### Strain engineering, activation and refactoring of silent BGC

In addition, complex BGC activations are also possible with our developed method. The activation of the putative safracin-associated BGC, named *xsc*, in *Xenorhabdus sp*. TS4 is such an example due to its bidirectional architecture (Fig. 2A I). In initial experiments, the cluster was amplified in three parts and assembled on expression vectors, which were subsequently transformed into *E*. *coli mtaA* in various combinations (Fig. S6). It was possible to produce different safracin derivatives with putative biosynthetic intermediates (Fig. S6 A-C). However, only small amounts of about 14 mg/L of SAC-B were produced (Fig 2 C). For this reason, we decided to use our developed method for targeted cluster refactoring. For activation of the *xsc* cluster (Fig. 2 A I), the modified strong inducible *vanR*/P_vanCC_ promoter was introduced upstream of *xscA* and a constitutive promoter (P_proC_) in the opposite direction upstream of *xscI* (Fig. 2 A IIa). In addition, the 5’-UTR between the RiboJ and the start codon was replaced with a translation enhancer sequence (g10T7 UTR (also abbreviated as T7g10), BBa_K1758100 (http://parts.igem.org/Part:BBa_K1758100)). Besides the expected main products SAC-A and SAC-B, some putative biosynthetic precursors could be identified, which mainly indicated a weak procession by the SAM dependent N-methyltransferase XscI and FAD-dependent monooxygenase XscJ. To overcome this, we replaced P_proC_ with a stronger constitutive promoter P_proD_ in front of *xscI*, resulting in strain IIb (Fig. 2 A). Furthermore, to gain control over the prodrug/drug formation and to obtain exclusively SAC-B derivatives, we replaced the promoter upstream of *xscK* encoding the peptidase with a modified *lacI*/P_tac_ IPTG inducible promoter while simultaneously deleting the putative resistance gene *xscH* [67]. The resulting strain III showed an exclusive and increased production of SAC-B in the range of 208±3.5 mg/L. To achieve an even higher level of SAC-B production, we also tried to increase the availability of the modified tyrosine precursor 3-hydroxy-5-methyl-*O*-methyltyrosine (Fig. 2 B) [68]. For this purpose, we introduced the biosynthetic genes *xscD*-*G* responsible for its formation on an L-arabinose inducible expression vector into strain III and used the empty vector as control. The introduction of the empty vector led to a drop in SAC-B production, which we assume was caused by stress from the added kanamycin. However, with the introduction of *xscD*-*G*, this effect could be compensated and a production titer of 336±26.6 mg/L could be achieved Fig. 2 C). Base peak chromatograms of the described strains can be found in Figure S8 A.

**Figure 2.**
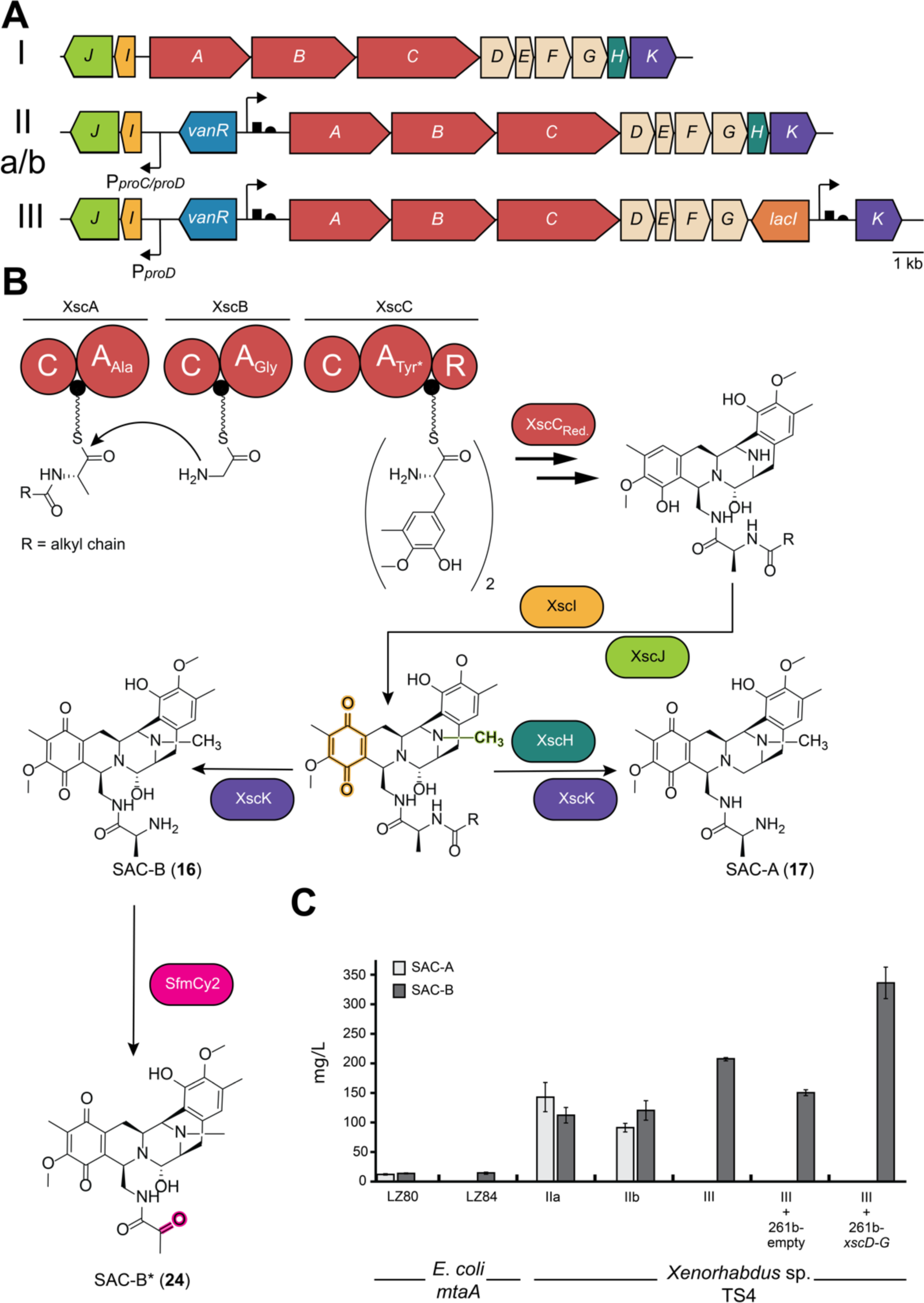
Safracin cluster activation and refactoring in *Xenorhabdus* sp. TS4. **A**) Constructed safracin producer strains, I represents the wildtype strain, IIa promoter exchange strain with P*_van_*/P*_proC_* insertion, IIb P*_van_*/P*_proD_* insertion and strain II with additional P_tac_ insertion upstream of *xscK* with *xscH* deletion. **B**) Putative biosynthesis of safracins; tailoring enzyme modifications are highlighted; Tyr* indicates the specificity of the XscC A domain for 3-hydroxy-5-methyl-o-methyltyrosine; “Unnatural” further modification of SAC-B by SfmCy resulting in **24**. **C**) Comparison of SAC-A and SAC-B production between the constructed strains. A: adenylation domain, C: condensation domain, R: reduction domain, PCP: peptidylcarrier protein (small black circle).

In addition to increasing the concentration of SAC-B, we also tested whether the product could be further functionalized. It was previously shown [36] that other “unnatural” safracin derivatives could be generated by adding modified precursors such as 3-hydroxy-5-methyl-O-ethyltyrosine. It was also shown previously [69] that functionalization of SAC-B by enzymes from the saframycin biosynthetic pathway of *S*. *lavendulae* was possible. We therefore introduced the codon-optimized *sfmO4* gene, which encodes a cytochrome P450 monooxygenase responsible for the introduction of an additional hydroxyl group. However, none of the previously described derivatives could be found, probably because the partnering reductase was missing (data not shown). In addition, we introduced the codon-optimized *sfmCy2* gene, encoding a secreted FAD-dependent oxidoreductase, by integration into the genome (Figure S7 B). SfmCy2 is responsible for the periplasmic oxidative deamination of the N-terminus [70]. After induction of all components in the production strain, we were able to detect a new mass of *m/z* = 522.223 [M+H-H_2_O]^+^ (**24**; Fig. 2B, S8 B). We hypothesize that this is the oxidatively deaminated derivative of SAC-B (**24**, Fig. 2B), which is supported by the MS-MS fragmentation pattern (Fig. S7 C). However, further studies are required to confirm its structure.

## Conclusions

In this work, we describe a simple and easy-to-use method for markerless genome editing in *Photorhabdus* and *Xenorhabdus*. To the best of our knowledge, this is the first description of CRISPR technology in the genus *Xenorhabdus*. The reduction to a single plasmid makes it much easier to work with non-laboratory organisms, which often show reduced growth with two resistance markers. The insertion of synthetic dsDNA fragments carrying both the repair template and the CRISPR target sequence makes vector assembly relatively simple. Furthermore, the easy transferability of the pSEVA vectors by conjugation, enabled by oriT, opens up the potential application to many different bacteria. In contrast to previously described genome editing methods, the process described here is straightforward and does not require the stepwise integration of one or more plasmids into the genome [33, 71]. From the transfer of the vector to the desired and resistance-free production strain, only four steps are required, which can be performed in six days (Fig. 3). On day seven, a new round of editing can be started, allowing rapid sequential editing. In “classical” BGC activation, R6K ori-based plasmids or a resistance cassette are often introduced in front of the desired gene, which has the disadvantage that iterative rounds of activation are not possible. [22, 32]. Due to the presence of the *sacB* gene on our vector and the fact that it is freely replicable, it can be easily cured by the addition of sucrose. Furthermore, our method is efficient for small and large deletions, as well as for smaller insertions, such as the insertion of inducible promoter sequences, without producing chimeric colonies (Table 1). We hypothesize that this is due to the use of two CRISPR sites per target. Targeting both ends of the sequence to be edited increases efficiency, as has been shown for *Streptomyces* [72]. Based on our technology, production titers of the high-value compound safracin B could be increased far beyond titers previously described [36]. Genomic integration of modifying enzymes also increased product variability and yielded a new, previously undescribed derivative **24**. The application of the single plasmid system and pSEVA based expression plasmids should therefore facilitate a variety of future studies in *Photorhabdus*, *Xenorhabdus*, as well as other bacteria, including regulatory network analysis, activation of silent natural product gene clusters, introduction of novel metabolic pathways, and design of optimized chassis strains for heterologous production of desired natural products.

**Figure 3.**
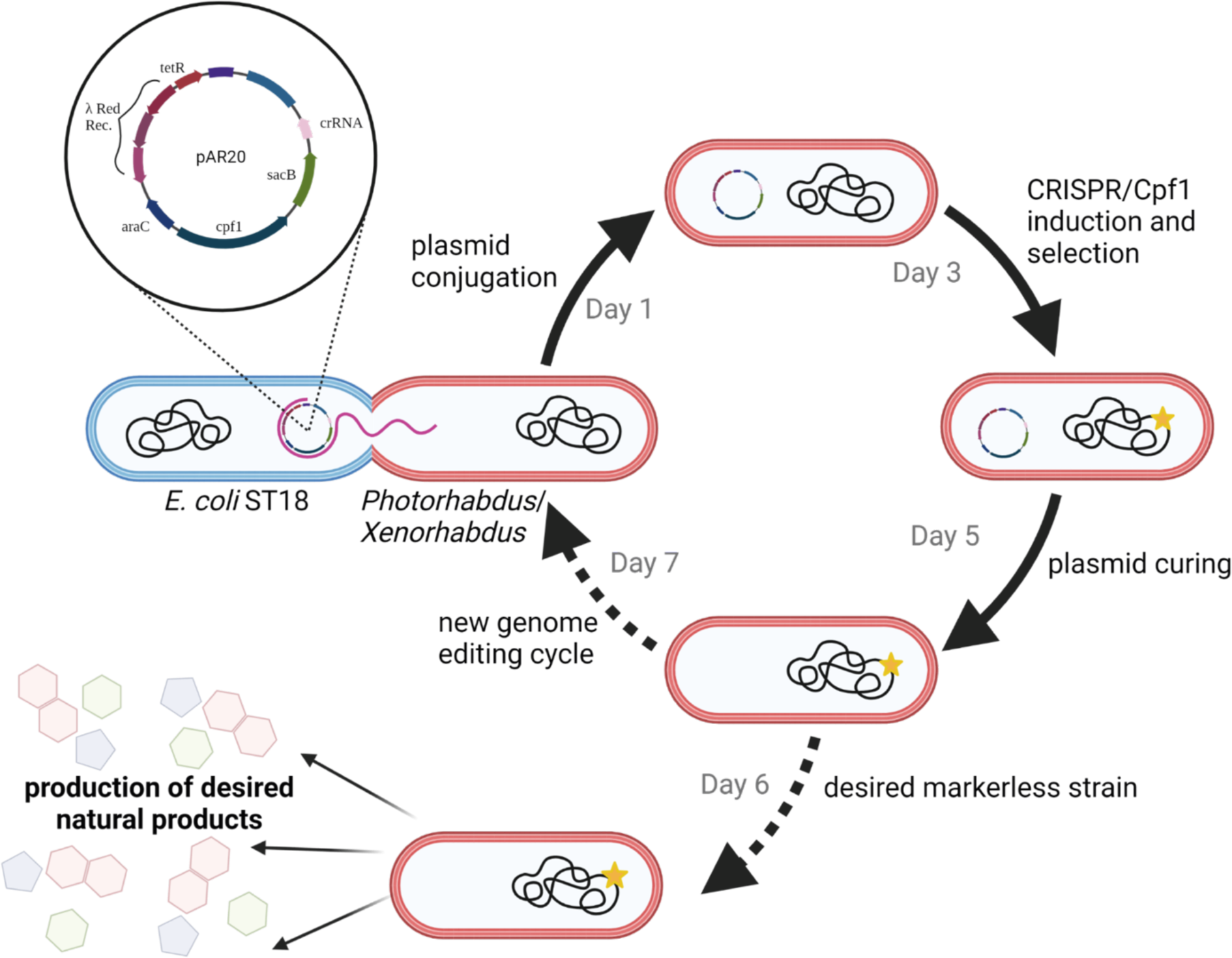
Overview of the developed single plasmid CRISPR-Cpf1 genome editing workflow. The approach starts with conjugation of the target specific pAR20 into the strain to be edited (day 1); selection of conjugates and induction of CRISPR/Cpf1 components followed by selection of correct edited strains (day 3); plasmid curing with sucrose (day 5); screening for new or known compounds (day 6); entering a new editing cycle (day 7). Figure created with BioRender.com.

## Acknowledgements

The authors are grateful to Peter Grün for help with the HPLC/MS analysis and maintenance of the analytical and preparative HPLC/MS systems. Furthermore, we thank Stefanie Lamberth for laboratory maintenance and preparation of large-scale cultures.

## Availability of data and materials

All data generated or analyzed during this study are included in this article and its supplementary information files.

## Author information

### Authors and Affiliations

Department of Natural Products in Organismic Interactions, Max-Planck Institute for Terrestrial Microbiology, 35043 Marburg, Germany

Alexander Rill, Helge B. Bode

Molecular Biotechnology, Department of Biosciences, Goethe University Frankfurt, 60438 Frankfurt am Main, Germany

Alexander Rill, Lei Zhao, Helge B. Bode

Department of Chemistry, Chemical Biology, Phillips University Marburg, 35043 Marburg, Germany

Alexander Rill, Helge B. Bode

Center for Synthetic Microbiology (SYNMIKRO), Phillips University Marburg, 35043 Marburg, Germany

Helge B. Bode

Senckenberg Gesellschaft für Naturforschung, 60325 Frankfurt, Germany

Helge B. Bode

State Key Laboratory of Bio-organic and Natural Products Chemistry, Shanghai Institute of Organic Chemistry, Chinese Academy of Sciences, 200032 Shanghai, China

Lei Zhao

### Contributions

Alexander Rill: method development, genetic construction, strain engineering data preparation and visualization, paper writing

Lei Zhao: vector construction for heterologous expression of *xsc* BGC in *E. coli*, isolation and structure elucidation of compound 20

Helge B. Bode: supervision, paper writing

### Corresponding author

Correspondence to Helge B. Bode

### Ethics approval and consent to participate

Not applicable.

### Consent for publication

Not applicable.

