## Additional File_1 for "Genetic Toolbox for *Photorhabdus* and *Xenorhabdus*: pSEVA based heterologous expression systems and CRISPR/Cpf1 based genome editing for rapid natural product profiling"

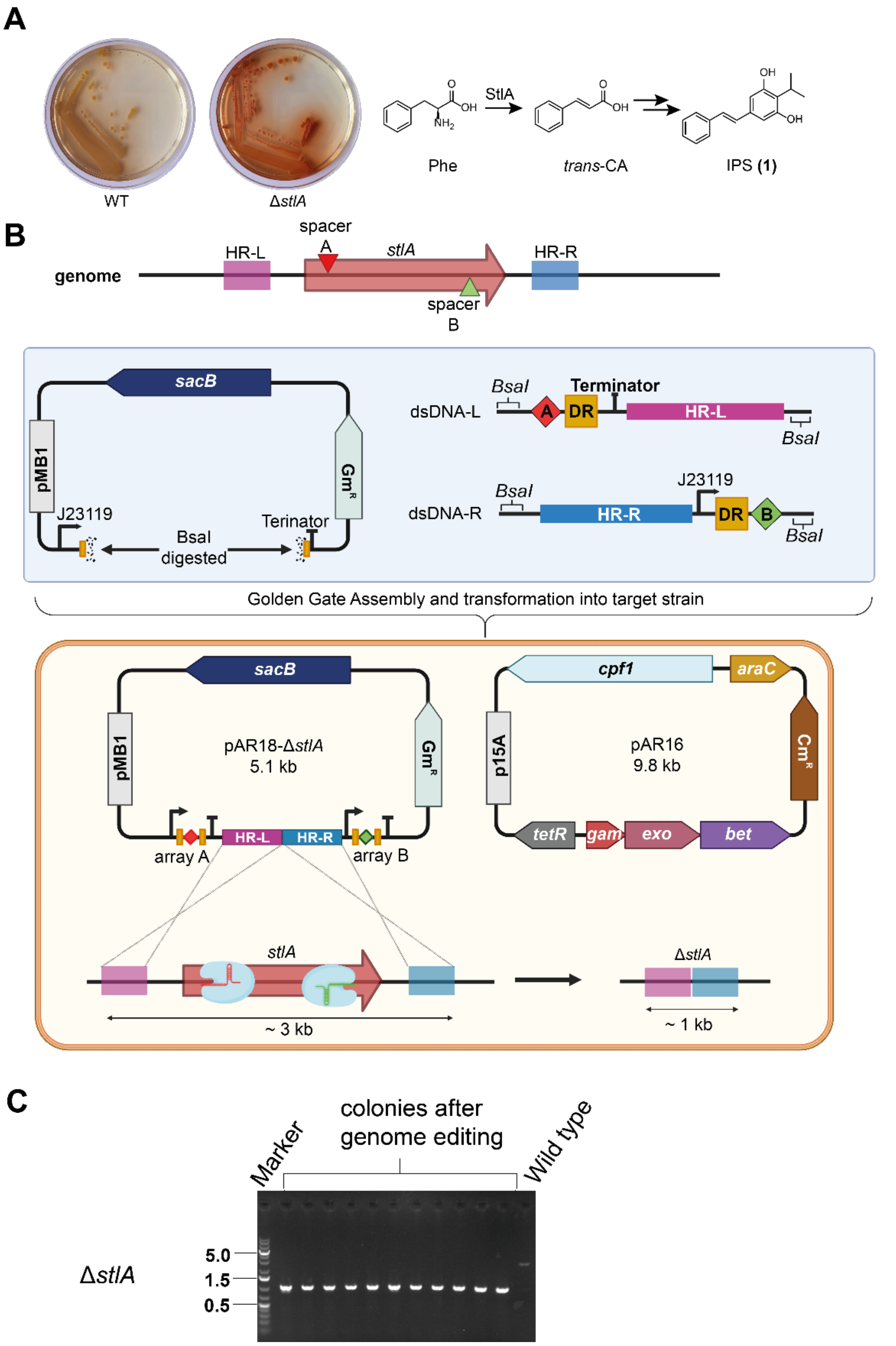


**Figure S1.** Design of Two plasmid based genome editing in *Photorhabdus* using *stlA* deletion as an example. A) Phenotypic comparison of wild type (WT) and Δ*stlA* deletion mutant; role of StlA in the biosynthesis of isopropylstilbene (1). B) Schematic representation of the selection of target sequence and homology arms; schematic cloning of synthetic dsDNA fragments harboring the repair template and target sequence into synthetic crRNA framework encoded on pAR18; detailed genotype of pAR18 are shown in Table S1. C) Gel electrophoresis image of deletion a mutants after colony PCR compared to the wild type. Taken from BioRender.com and further modified.

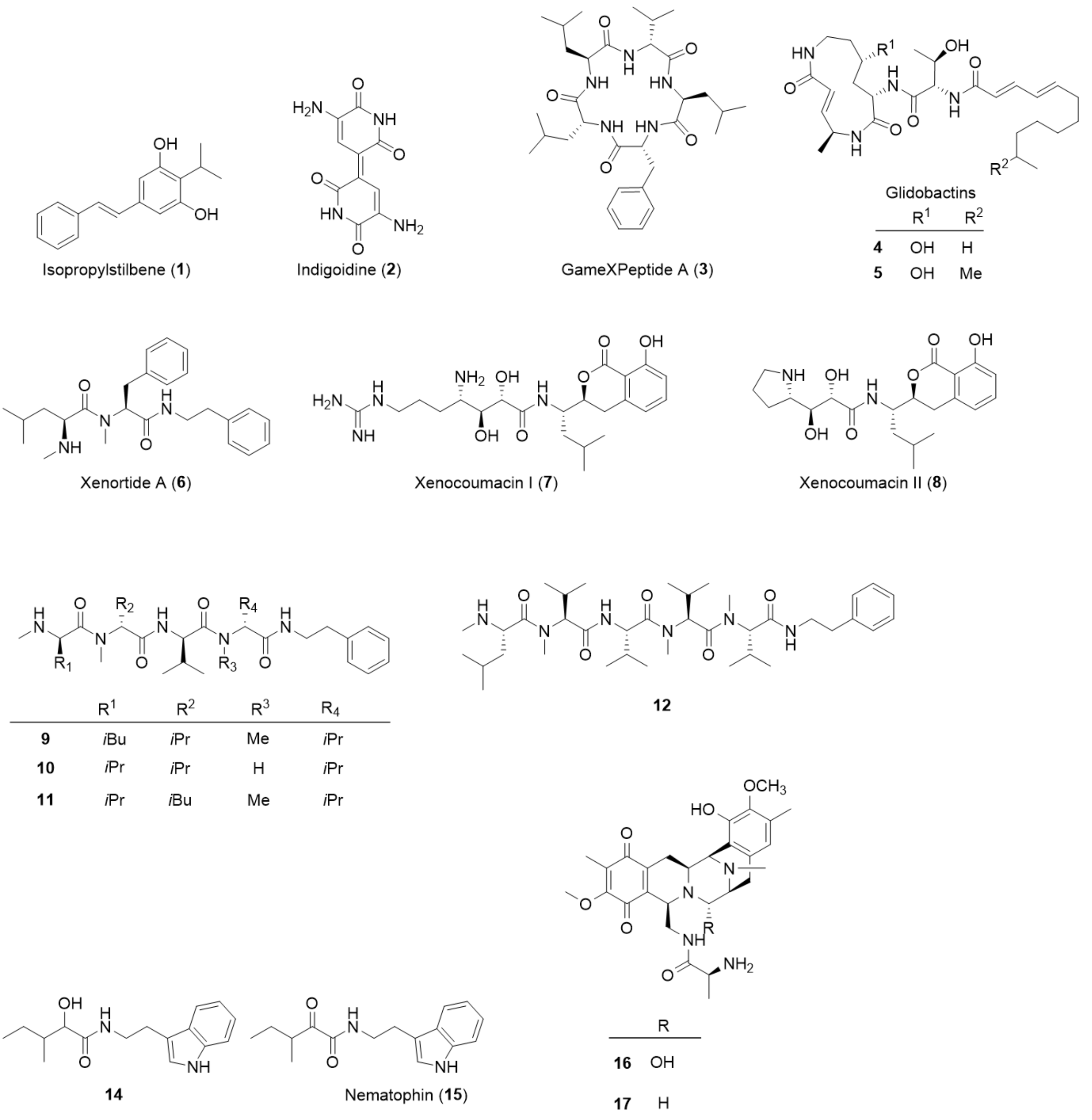


**Figure S2. Structures of described compounds.** Compound **13** is described in [1] and compound **15** in [2] while compound **14** was confirmed by sum formula prediction as partly reduced derivative of **15**.

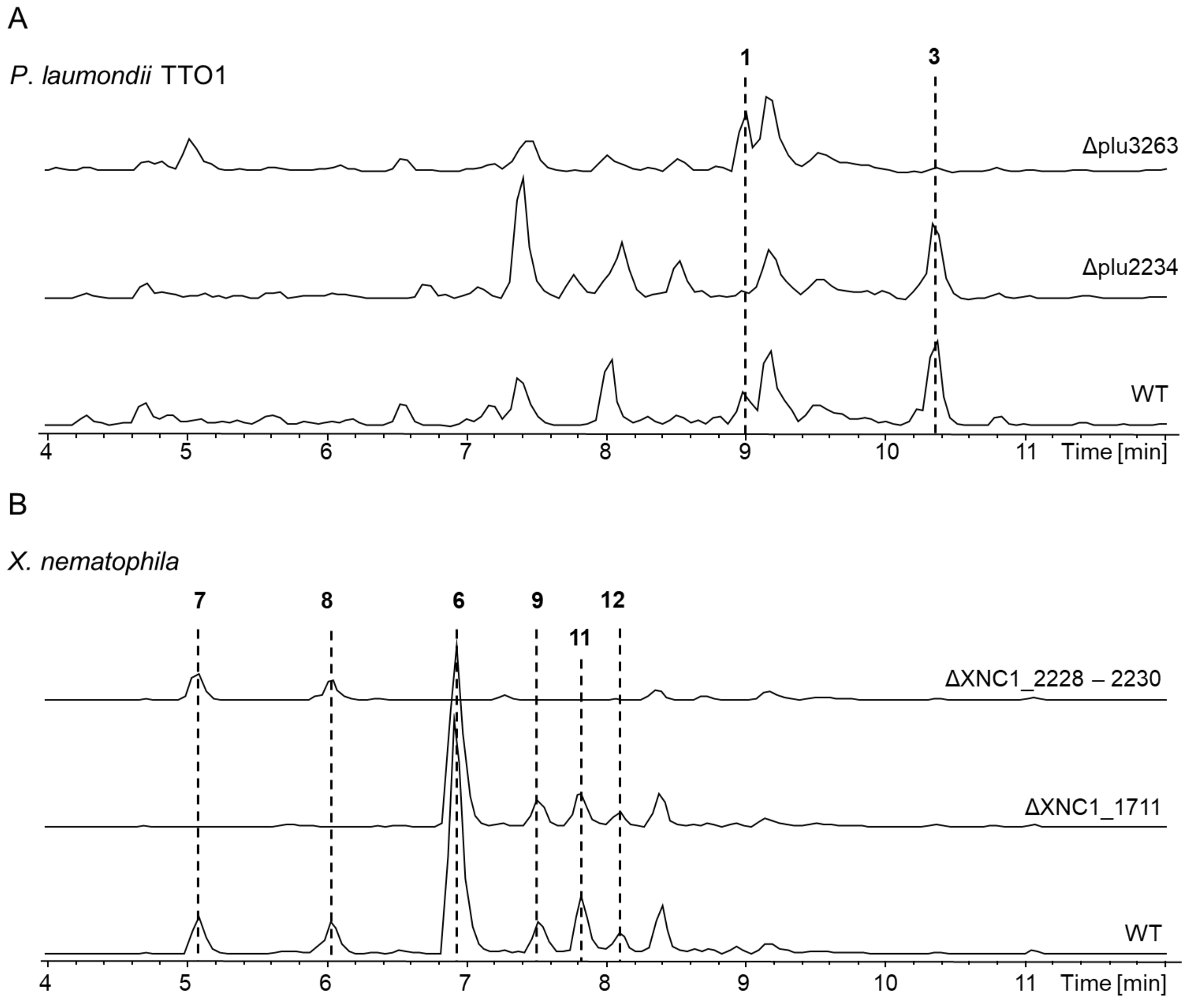


**Figure S3. Overview of MS spectra of deletion mutants.** A) Base peak chromatograms (BPC) of *P. laumondii* TTO1 wild type and deletion mutants. B) BPCs of *X. nematophila* wild type and deletion mutants.

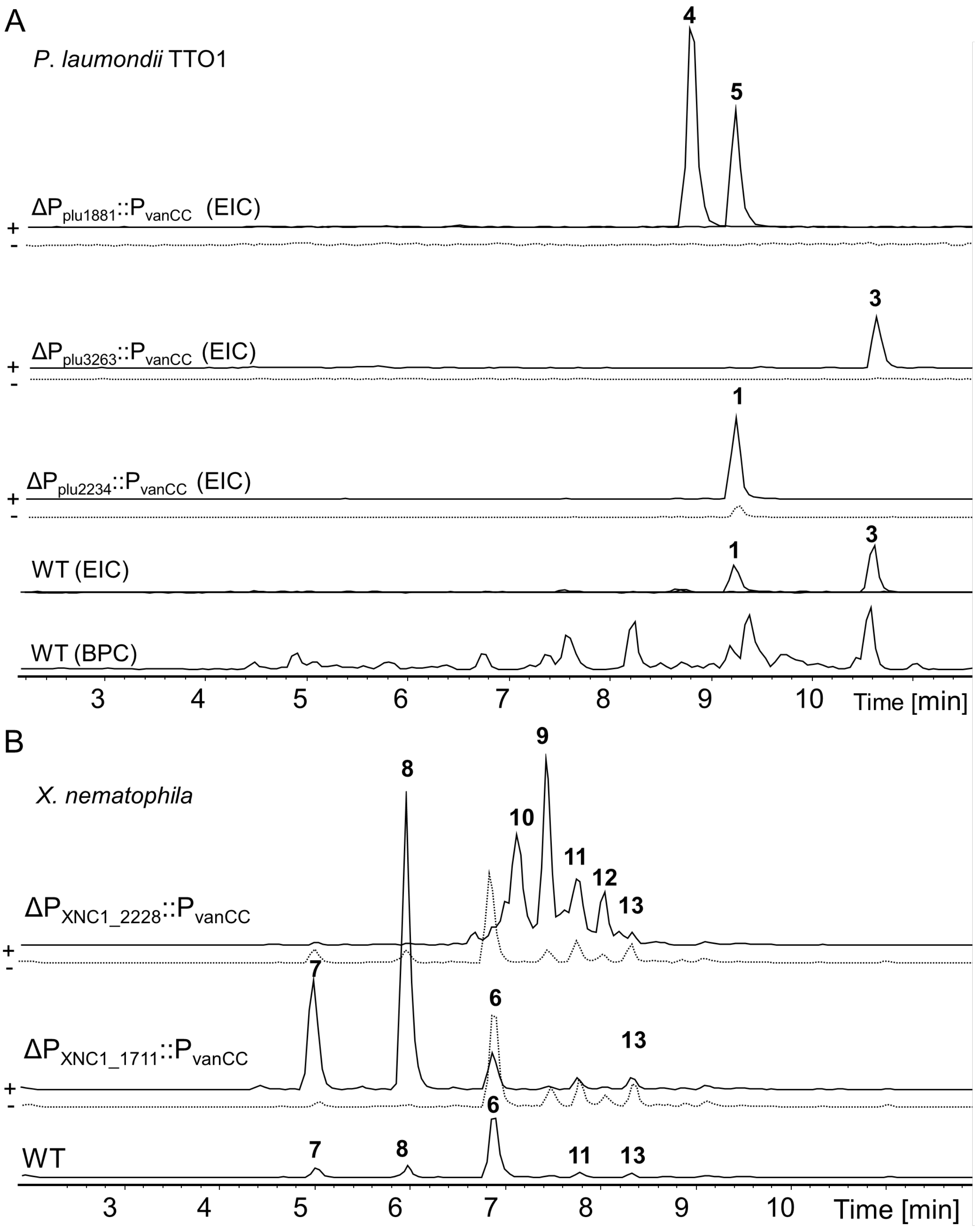


**Figure S4. Overview of MS spectra of induced (solid line) and non-induced (dashed line) cultures after successful promoter exchanges**. A) Chromatograms of *P. laumondii* TTO1 and edited mutants; wild type (WT) BPC and for better visualization extracted ion chromatograms (EICs) representing major derivatives of associated NPs. B) Chromatograms of *Xenorhabdus nematophila* wild type and genome edited mutants. (NPs mentioned are shown in Figure S1).

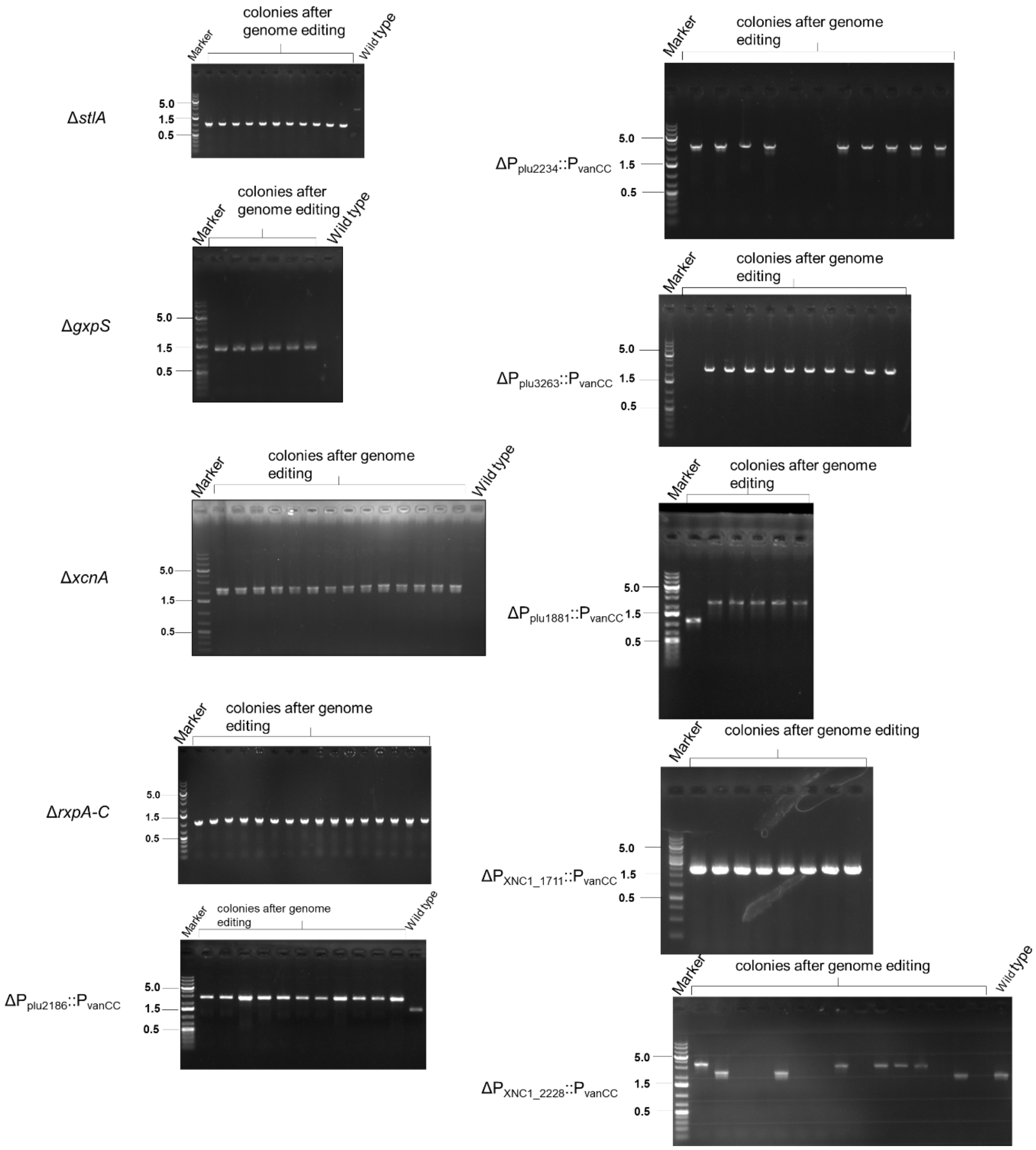


**Figure S5. Gel electrophoresis images of deletion and promoter exchange mutants after colony PCR.**

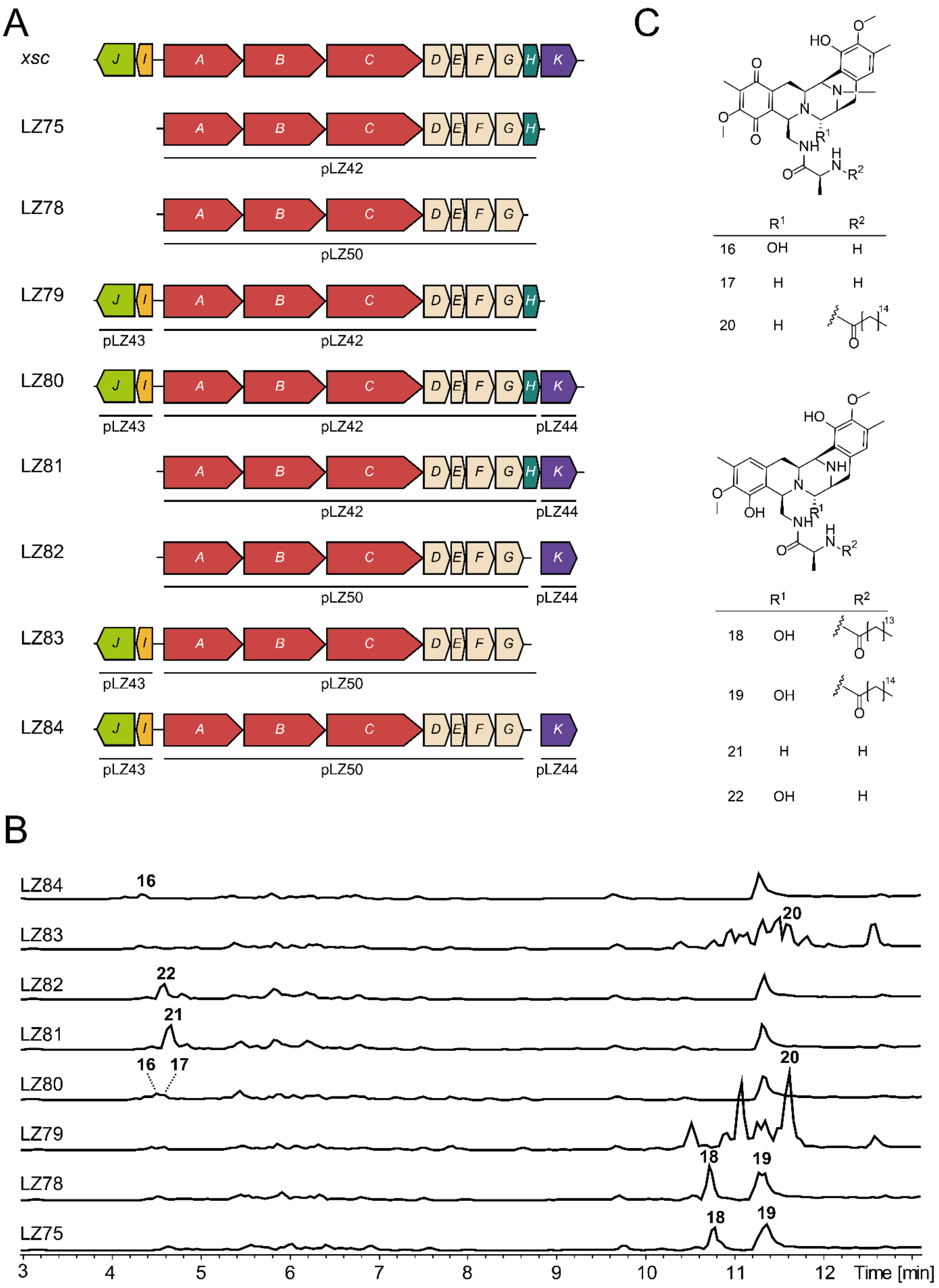


**Figure S6. Heterologous production of various combinations of the putative safracin BGC *xsc* from *Xenorhabdus* sp. TS4 in *E. coli mtaA*.** A) overview about the different plasmid combinations. B). Base peak chromatograms of safracin derivatives producing *E. coli* mtaA strains. C) Putative structures of safracin biosynthesis intermediates and final products SAC-A and SAC-B.

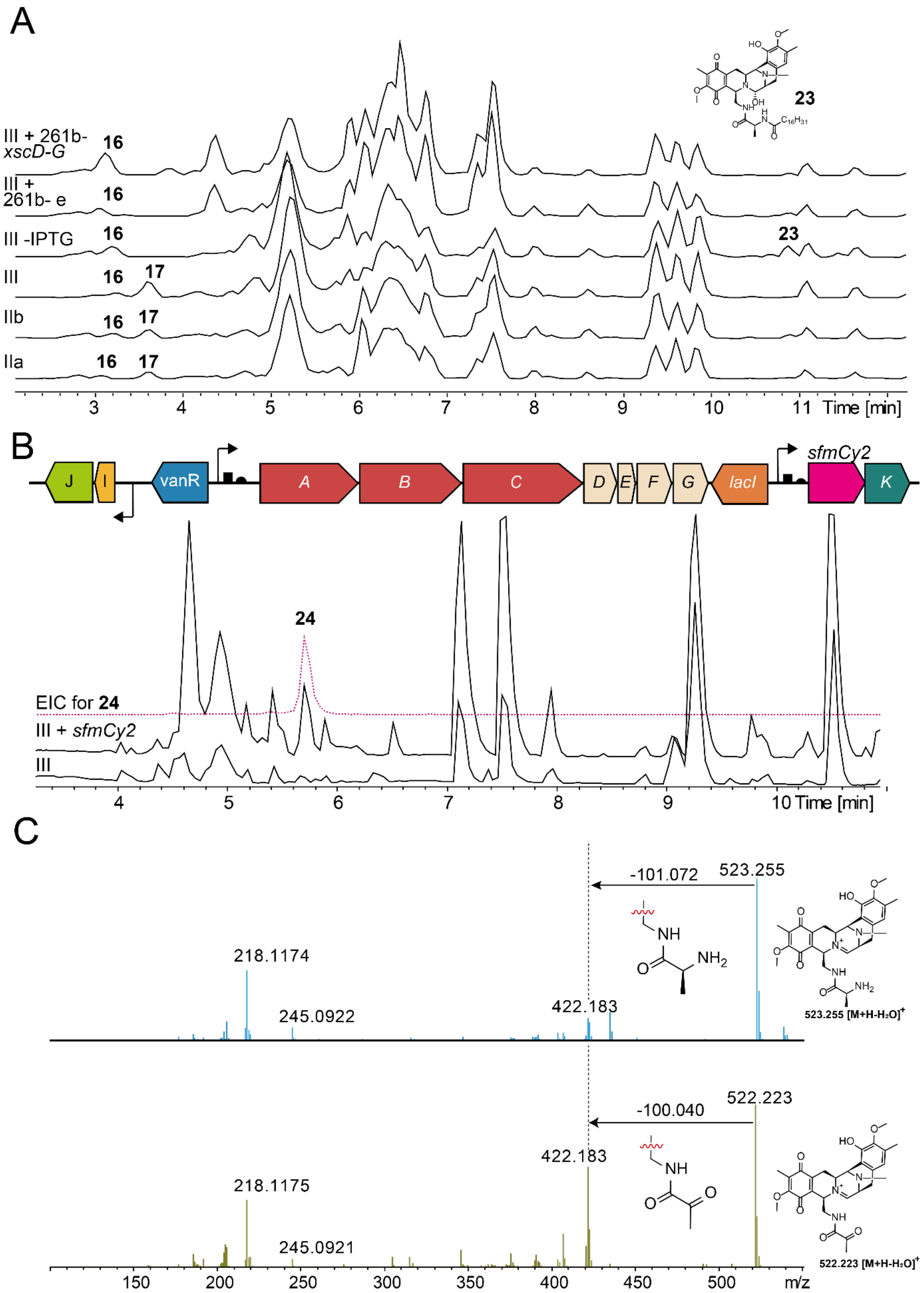


**Figure S7. MS analysis of *Xenorhabdus* sp. TS4 extracts after *xsc* BGC induction.** A) Base peak chromatograms of strains described in Figure 2, putative structure of the major acyl-derivative **23** of SAC-B in a production culture without *xscK* induction. B) genomic integration of *sfmCy2* upstream of *xscK*, both under the control of IPTG inducible promoter $P_{tac}$. Base preak chromatogram of selected strain with *sfmCy2* integration compared to strain III. C) MS2 fragmentation pattern of **16** compared to **24**. Mass loss and putative structure are indicated with arrows.

**Table S1.** Strains and plasmids used in this study.

| Strains and plasmids | Characteristics | Source/Reference |
|---|---|---|
| Strain | | |
| *E. coli* ST18 | pro thi hsdR$^+$ Tp$^r$ Sm$^r$; chromosome::RP4-2 Tc::Mu-Kan::Tn7/λpir; Δ*hemA* | [3] |
| *E. coli* DH10B | F$^-$*mcr*A Δ(*mrr-hsd*RMS-*mcr*BC) φ80*lac*ZΔM15 Δ*lac*X74 *rec*A1 *end*A1 *ara*D139 Δ(*ara-leu*)7697 *gal*U *gal*K λ$^-$*rps*L(Str$^R$) *nup*G | Invitrogen |
| *Photorhabdus laumondii* subsp. *laumondii* TTO1 | Wild type | |
| *Xenorhabdus nematophila* ATCC 19061 | Wild type | |
| ARs137 | *P. laumondii* TTO1 Δplu2234 | This study |
| ARs139 | *P. laumondii* TTO1 Δplu3263 | This study |
| ARs140 | *P. laumondii* TTO1 $\Delta P_{plu2186}$::$P_{vanCC}$ | This study |
| ARs141 | *P. laumondii* TTO1 $\Delta P_{plu2234}$::$P_{vanCC}$ | This study |
| ARs142 | *P. laumondii* TTO1 $\Delta P_{plu3263}$::$P_{vanCC}$ | This study |
| ARs143 | *P. laumondii* TTO1 $\Delta P_{plu1881}$::$P_{vanCC}$ | This study |
| ARs157 | *X. nematophila* ΔXNC1_2228 – 2230 | This study |
| ARs158 | *X. nematophila* ΔXNC1_1711 | This study |
| ARs159 | *X. nematophila* $\Delta P_{XNC1\_2228}$::$P_{vanCC}$ | This study |
| ARs160 | *X. nematophila* $\Delta P_{XNC1\_1711}$::$P_{vanCC}$ | This study |
| LZ75 | *E. coli mtaA* + pLZ42 | This study |
| LZ78 | *E. coli mtaA* + pLZ50 | This study |
| LZ79 | *E. coli mtaA* + pLZ42 + pLZ43 | This study |
| LZ80 | *E. coli mtaA* + pLZ42 + pLZ43 + pLZ44 | This study |
| LZ81 | *E. coli mtaA* + pLZ42 + pLZ44 | This study |
| LZ82 | *E. coli mtaA* + pLZ50 + pLZ44 | This study |
| LZ83 | *E. coli mtaA* + pLZ50 + pLZ43 | This study |
| LZ84 | *E. coli mtaA* + pLZ50 + pLZ43 + pLZ44 | This study |
| ARs337 (strain I) | *Xenorhabdus* sp. TS4 $\Delta P_{XETS4V2\_00440}$::$P_{vanCC}$ $\Delta P_{XETS4V2\_00445}$::proC | This study |
| ARs337 (strain II) | *Xenorhabdus* sp. TS4 $\Delta P_{XETS4V2\_00440}$::$P_{vanCC}$ $\Delta P_{XETS4V2\_00445}$::proD | This study |
| ARs337 (strain III) | *Xenorhabdus* sp. TS4 $\Delta P_{XETS4V2\_00440}$::$P_{vanCC}$ $\Delta P_{XETS4V2\_00445}$::proD ΔXETS4V2_00405 $\Delta P_{XETS4V2\_00400}$::$P_{tac}$ | This study |
| ARs337 + 261e | ARs337 + 261e | This study |
| ARs337 + 261-*xscD-G* | ARs337 + 261-*xscD-G* | This study |
| Plasmid | | |
| pTargetF | pMB1 Spec$^R$ sgRNA-cadA | pTargetF was a gift from Sheng |

| | | |
|---|---|---|
| | | Yang (Addgene plasmid # 62226)[4] |
| p46Cpf1-OP2 | repA101(Ts) *cat tetR* $P_{tet}$-Red *araC* $P_{Bad}$ codon optimized FnCas12a | p46Cpf1-OP2 was a gift from Qiong Wu (Addgene plasmid # 98592)[5] |
| pAR14 | pMB1 $Spec^R$ *sacB* | This study |
| pAR15 | pMB1 $Gm^R$ *sacB* | This study |
| pAR16 | p15A cat tetR $P_{tet}$-Red araC $P_{Bad}$ codon optimized FnCas12a | This study |
| pAR18 | pMB1 $Gm^R$ *sacB* crRNA framework | This study |
| pSEVA231 | pBBR1 ori $Km^R$ oriT | [6] |
| pAR20 | pSEVA231 *tetR* $P_{tet}$-λRed *araC* $P_{Bad}$ codon optimized FnCas12a *sacB* crRNA framework | This study |
| pSEVA261 | p15A ori $Km^R$ oriT | [6] |
| pSEVA221 | RK2 ori $Km^R$ oriT | [6] |
| pSEVA341 | pRO1600_ColE1 ori $Cm^R$ oriT | [6] |
| pSEVA351 | RSF1010 ori $Cm^R$ oriT | [6] |
| pSEVA621 | RK2 ori $Gm^R$ oriT | [6] |
| pSEVA631 | pBBR1 ori $Gm^R$ oriT | [6] |
| pSEVA661 | p15A ori $Gm^R$ oriT | [6] |
| pSEVA221b-Neon | pSEVA221 $araC^{AM}$ $P_{Bad}$ RiboJ B0064-RBS mNeonGreen CDS | This study |
| pSEVA231b-Neon | pSEVA231 $araC^{AM}$ $P_{Bad}$ RiboJ B0064-RBS mNeonGreen CDS | This study |
| pSEVA261b-Neon | pSEVA261 $araC^{AM}$ $P_{Bad}$ RiboJ B0064-RBS mNeonGreen CDS | This study |
| pSEVA221v-Neon | pSEVA221 *vanR* $P_{vanCC}$ RiboJ B0064-RBS mNeonGreen CDS | This study |
| pSEVA231v-Neon | pSEVA231 *vanR* $P_{vanCC}$ RiboJ B0064-RBS mNeonGreen CDS | This study |
| pSEVA261v-Neon | pSEVA261 *vanR* $P_{vanCC}$ RiboJ B0064-RBS mNeonGreen CDS | This study |
| pSEVA221t-Neon | pSEVA221 *lacI* $P_{tac}$ RiboJ B0064-RBS mNeonGreen CDS | This study |
| pSEVA231t-Neon | pSEVA231 *lacI* $P_{tac}$ RiboJ B0064-RBS mNeonGreen CDS | This study |
| pSEVA261t-Neon | pSEVA261 *lacI* $P_{tac}$ RiboJ B0064-RBS mNeonGreen CDS | This study |
| pSEVA621b-Neon | pSEVA621 $araC^{AM}$ $P_{Bad}$ RiboJ B0064-RBS mNeonGreen CDS | This study |
| pSEVA631b-Neon | pSEVA631 $araC^{AM}$ $P_{Bad}$ RiboJ B0064-RBS mNeonGreen CDS | This study |
| pSEVA661b-Neon | pSEVA661 $araC^{AM}$ $P_{Bad}$ RiboJ B0064-RBS mNeonGreen CDS | This study |
| pSEVA621v-Neon | pSEVA621 *vanR* $P_{vanCC}$ RiboJ B0064-RBS mNeonGreen CDS | This study |
| pSEVA631v-Neon | pSEVA631 *vanR* $P_{vanCC}$ RiboJ B0064-RBS mNeonGreen CDS | This study |

| | | |
|---|---|---|
| pSEVA661v-Neon | pSEVA661 *vanR* $P_{vanCC}$ RiboJ B0064-RBS mNeonGreen CDS | This study |
| pSEVA621t-Neon | pSEVA621 *lacI* $P_{tac}$ RiboJ B0064-RBS mNeonGreen CDS | This study |
| pSEVA631t-Neon | pSEVA631 *lacI* $P_{tac}$ RiboJ B0064-RBS mNeonGreen CDS | This study |
| pSEVA661t-Neon | pSEVA661 *lacI* $P_{tac}$ RiboJ B0064-RBS mNeonGreen CDS | This study |
| pSEVA341b-Neon | pSEVA341 $araC^{AM}$ $P_{Bad}$ RiboJ B0064-RBS mNeonGreen CDS | This study |
| pSEVA351b-Neon | pSEVA351 $araC^{AM}$ $P_{Bad}$ RiboJ B0064-RBS mNeonGreen CDS | This study |
| pSEVA341v-Neon | pSEVA341 *vanR* $P_{vanCC}$ RiboJ B0064-RBS mNeonGreen CDS | This study |
| pSEVA351v-Neon | pSEVA351 *vanR* $P_{vanCC}$ RiboJ B0064-RBS mNeonGreen CDS | This study |
| pSEVA341t-Neon | pSEVA341 *lacI* $P_{tac}$ RiboJ B0064-RBS mNeonGreen CDS | This study |
| pSEVA351t-Neon | pSEVA351 *lacI* $P_{tac}$ RiboJ B0064-RBS mNeonGreen CDS | This study |
| pSEVA341c-Neon | pSEVA341 *cymR* $P_{cymRC}$ RiboJ B0064-RBS mNeonGreen CDS | This study |
| pSEVA351c-Neon | pSEVA351 *cymR* $P_{cymRC}$ RiboJ B0064-RBS mNeonGreen CDS | This study |
| pSEVA221b | pSEVA221 $araC^{AM}$ $P_{Bad}$ RiboJ B0064-RBS | This study |
| pSEVA231b | pSEVA231 $araC^{AM}$ $P_{Bad}$ RiboJ B0064-RBS | This study |
| pSEVA261b | pSEVA261 $araC^{AM}$ $P_{Bad}$ RiboJ B0064-RBS | This study |
| pSEVA221v | pSEVA221 *vanR* $P_{vanCC}$ RiboJ B0064-RBS | This study |
| pSEVA231v | pSEVA231 *vanR* $P_{vanCC}$ RiboJ B0064-RBS | This study |
| pSEVA261v | pSEVA261 *vanR* $P_{vanCC}$ RiboJ B0064-RBS | This study |
| pSEVA221t | pSEVA221 *lacI* $P_{tac}$ RiboJ B0064-RBS | This study |
| pSEVA231t | pSEVA231 *lacI* $P_{tac}$ RiboJ B0064-RBS | This study |
| pSEVA261t | pSEVA261 *lacI* $P_{tac}$ RiboJ B0064-RBS | This study |
| pSEVA621b | pSEVA621 $araC^{AM}$ $P_{Bad}$ RiboJ B0064-RBS | This study |
| pSEVA631b | pSEVA631 $araC^{AM}$ $P_{Bad}$ RiboJ B0064-RBS | This study |
| pSEVA661b | pSEVA661 $araC^{AM}$ $P_{Bad}$ RiboJ B0064-RBS | This study |
| pSEVA621v | pSEVA621 *vanR* $P_{vanCC}$ RiboJ B0064-RBS | This study |
| pSEVA631v | pSEVA631 *vanR* $P_{vanCC}$ RiboJ B0064-RBS | This study |
| pSEVA661v | pSEVA661 *vanR* $P_{vanCC}$ RiboJ B0064-RBS | This study |
| pSEVA621t | pSEVA621 *lacI* $P_{tac}$ RiboJ B0064-RBS | This study |
| pSEVA631t | pSEVA631 *lacI* $P_{tac}$ RiboJ B0064-RBS | This study |
| pSEVA661t | pSEVA661 *lacI* $P_{tac}$ RiboJ B0064-RBS | This study |
| pSEVA341b | pSEVA341 $araC^{AM}$ $P_{Bad}$ RiboJ B0064-RBS | This study |
| pSEVA351b | pSEVA351 $araC^{AM}$ $P_{Bad}$ RiboJ B0064-RBS | This study |
| pSEVA341v | pSEVA341 *vanR* $P_{vanCC}$ RiboJ B0064-RBS | This study |
| pSEVA351v | pSEVA351 *vanR* $P_{vanCC}$ RiboJ B0064-RBS | This study |
| pSEVA341t | pSEVA341 *lacI* $P_{tac}$ RiboJ B0064-RBS | This study |
| pSEVA351t | pSEVA351 *lacI* $P_{tac}$ RiboJ B0064-RBS | This study |
| pSEVA341c | pSEVA341 *cymR* $P_{cymRC}$ RiboJ B0064-RBS | This study |
| pSEVA351c | pSEVA351 *cymR* $P_{cymRC}$ RiboJ B0064-RBS | This study |
| pCOLA | ColE1 ori $Km^R$ araC $P_{Bad}$ | [7] |
| pACYC | p15A pri $Cm^R$ araC $P_{Bad}$ | [7] |
| pCDF | CloDF13 $Spec^R$ araC $P_{Bad}$ | [7] |

| | | |
|---|---|---|
| pLZ42 | ColE1 ori $Km^R$ araC $P_{Bad}$-*xscA-H* | This study |
| pLZ43 | p15A pri $Cm^R$ araC $P_{Bad}$-*xscIJ* | This study |
| pLZ44 | CloDF13 $Spec^R$ araC $P_{Bad}$-*xscK* | This study |
| pLZ50 | ColE1 ori $Km^R$ araC $P_{Bad}$-*xscA-G* | This study |
| pSEVA261b-*xscD-G* | pSEVA261b- *xscD-G* | This study |

**Table S3.** Proteins of the *xsc* cluster in *Xenorhabdus* sp. TS4, their proposed function, protein size and closest homologues.

| Protein | NCBI Reference Sequence | Size (aa) | Proposed function |
|---|---|---|---|
| XscA | Xets_RS15840 | 1055 | non-ribosomal peptide synthetase |
| XscB | Xets_RS15835 | 1073 | non-ribosomal peptide synthetase |
| XscC | Xets_RS15830 | 1444 | non-ribosomal peptide synthetase |
| XscD | Xets_RS15825 | 352 | putative hydroxylase |
| XscE | Xets_RS15820 | 66 | MbtH family NRPS accessory protein |
| XscF | Xets_RS15815 | 355 | O-methyltransferase |
| XscG | Xets_RS15810 | 343 | O-methyltransferase |
| XscH | Xets_RS15805 | 180 | dihydrofolate reductase/ dehydratase |
| XscI | Xets_RS15845 | 220 | SAM-dependent methyltransferase |
| XscJ | Xets_RS15850 | 501 | FAD-dependent monooxygenase |
| XscK | Xets_RS24810 | 512 | metallopeptidase |

**Table S4.** $^{1}H$ (600 MHz) and $^{13}C$ (150 MHz) NMR data of compound **20** in DMSO-$d_6$ ($\delta$ in ppm)

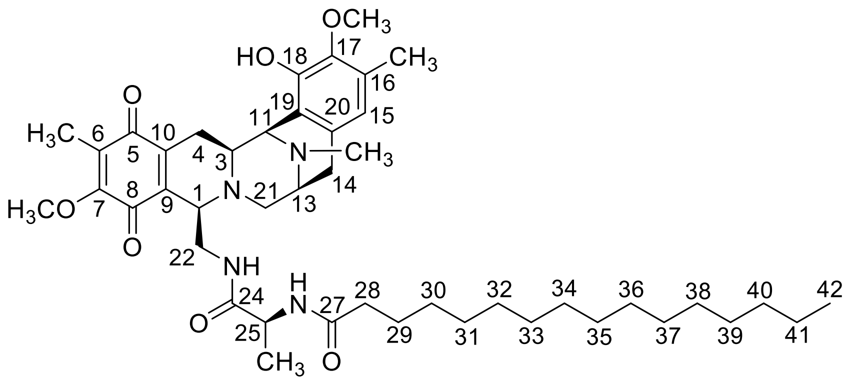

| position | $\delta_C$[a], type | $\delta_H$ ($J$ in Hz) |
|---|---|---|
| 1 | 58.1, CH | 3.29, m |
| 3 | 57.5, CH | 2.61, m |
| 4 | 24.7, $CH_2$ | 2.78, m; 1.70, m |
| 5 | 185.5, C | |
| 6 | 126.5, C | |
| 6-$CH_3$ | 8.2, $CH_3$ | 1.78, s |
| 7 | 155.3, C | |
| 7-$OCH_3$ | 59.8, $CH_3$ | 3.84, s |
| 8 | not detected, C | |
| 9 | 154.1, C | |
| 10 | 135.9, C | |
| 11 | 55.8, CH | 3.96, brs |
| 12-$NCH_3$ | 40.9, $CH_3$ | 2.12, s |
| 13 | 51.9, CH | 2.99, m |
| 14 | 25.7, $CH_2$ | 2.84, m; 2.55, d (17.5) |
| 15 | 119.2, CH | 6.38, s |
| 16 | 127.8, C | |
| 16-$CH_3$ | 15.2, $CH_3$ | 2.13, s |
| 17 | 142.5, C | |
| 17-$OCH_3$ | 59.3, $CH_3$ | 3.52, s |
| 18 | not detected, C | |
| 19 | 118.3, C | |
| 20 | 131.8, C | |
| 21 | 58.7, $CH_2$ | 3.08, d (8.7); 2.61, m |
| 22 | 39.4, $CH_2$ | 3.41, m; 3.28, m |
| 22-NH | | 7.41, d (6.8) |
| 24 | 171.6, C | |
| 25 | 47.9, CH | 3.41, m |
| 25-$CH_3$ | 17.4, $CH_3$ | 0.67, d (5.4) |
| 25-NH | | 6.19, m |
| 27 | 174.2, C | |
| 28 | 33.6, $CH_2$ | 2.16, m |
| 29 | 24.2, $CH_2$ | 1.49, m |
| 30-39 | 27.4-29.8, $CH_2$ | 1.25, m |
| 40 | 30.9, $CH_2$ | 1.25, m |
| 41 | 21.8, $CH_2$ | 1.26, m |
| 42 | 13.7, $CH_3$ | 0.85, t (7.0) |

[a]Due to weak $^{13}C$ signals, the data were deduced by the HSQC and HMBC spectra.

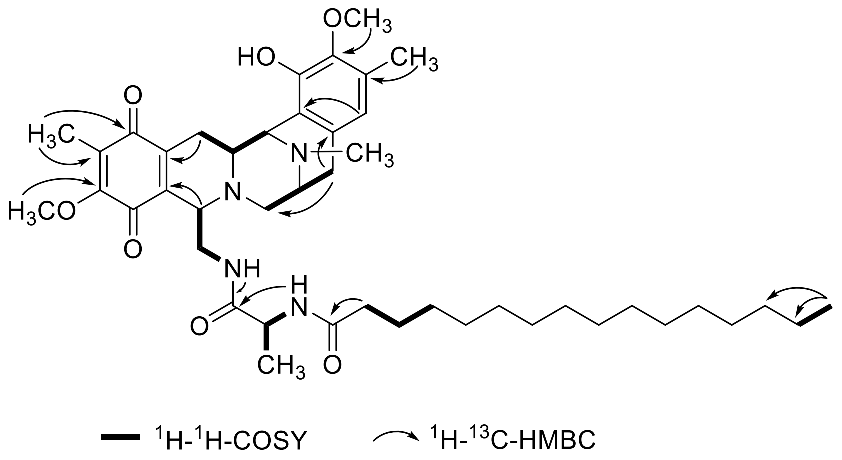


**Figure S8.** Key COSY and HMBC correlations of compound **20**.

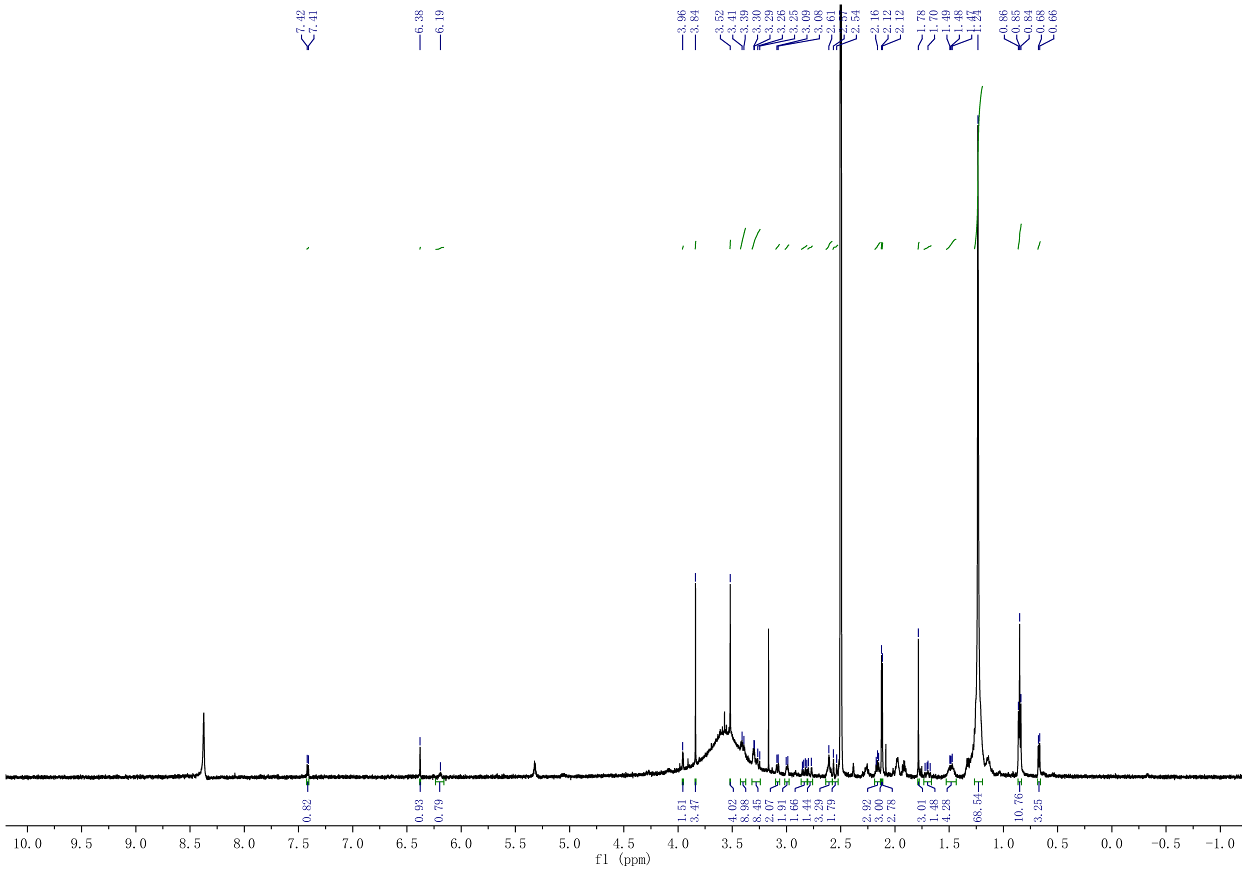


**Figure S9.** $^1$H NMR (600 MHz) spectrum of compound **20** in DMSO-$d_6$.

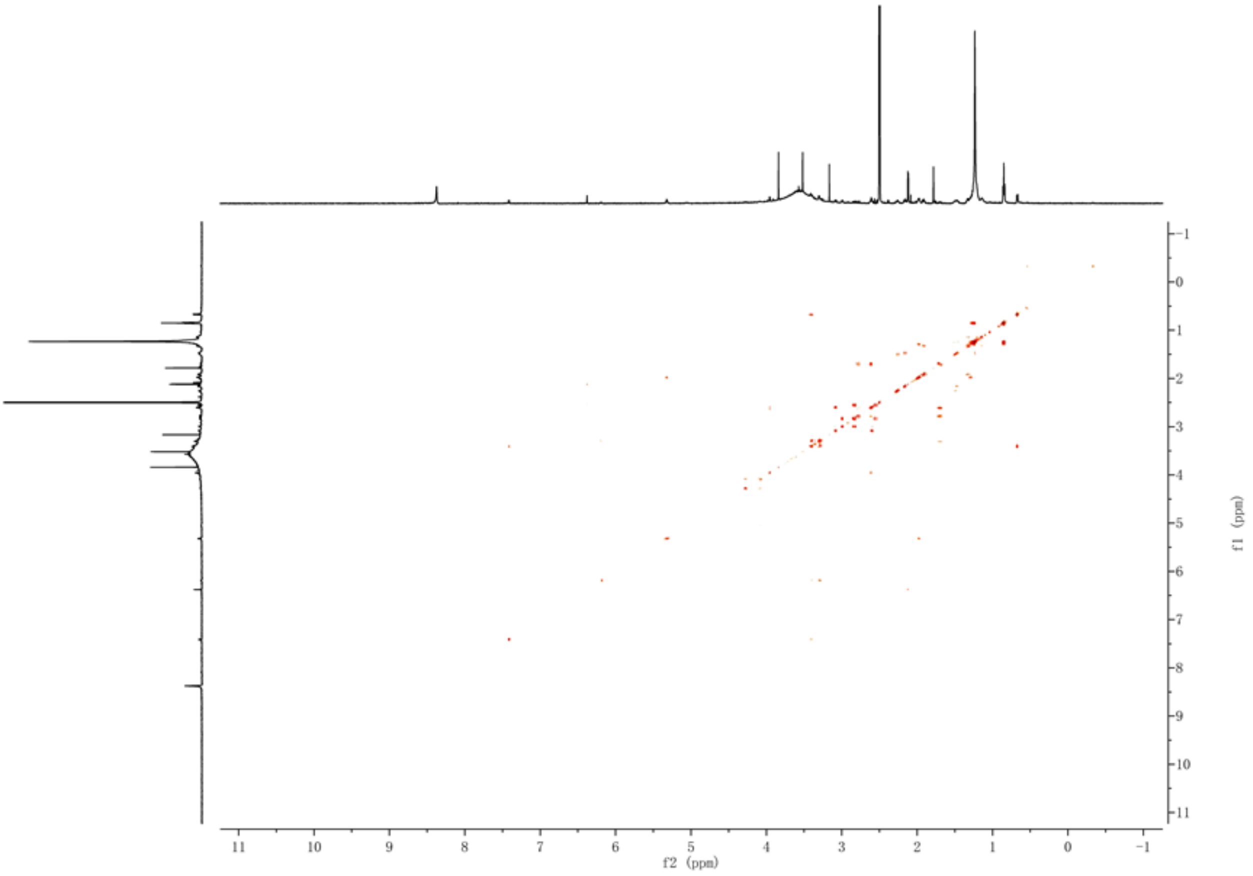


**Figure S10.** COSY spectrum of compound **20** in DMSO-$d_6$.

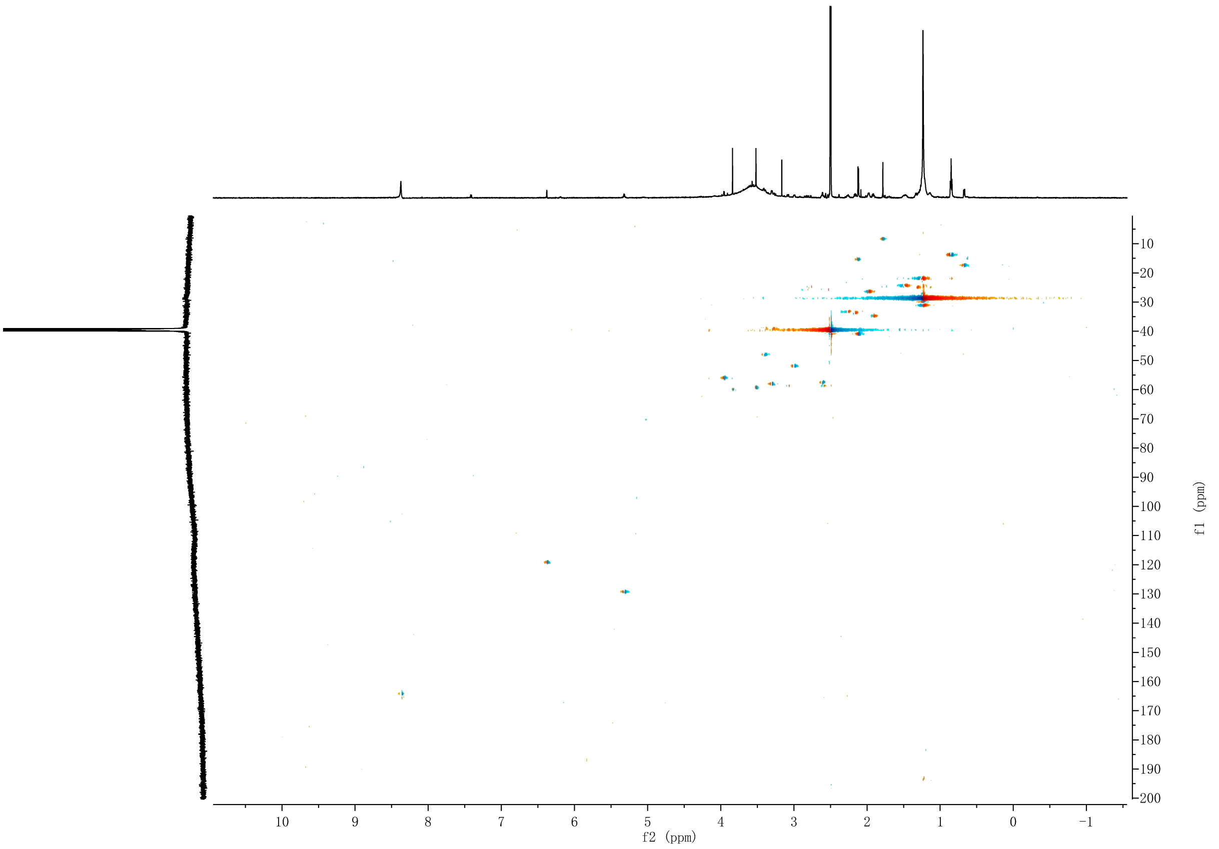


**Figure S11.** HSQC spectrum of compound **20** in DMSO-$d_6$.

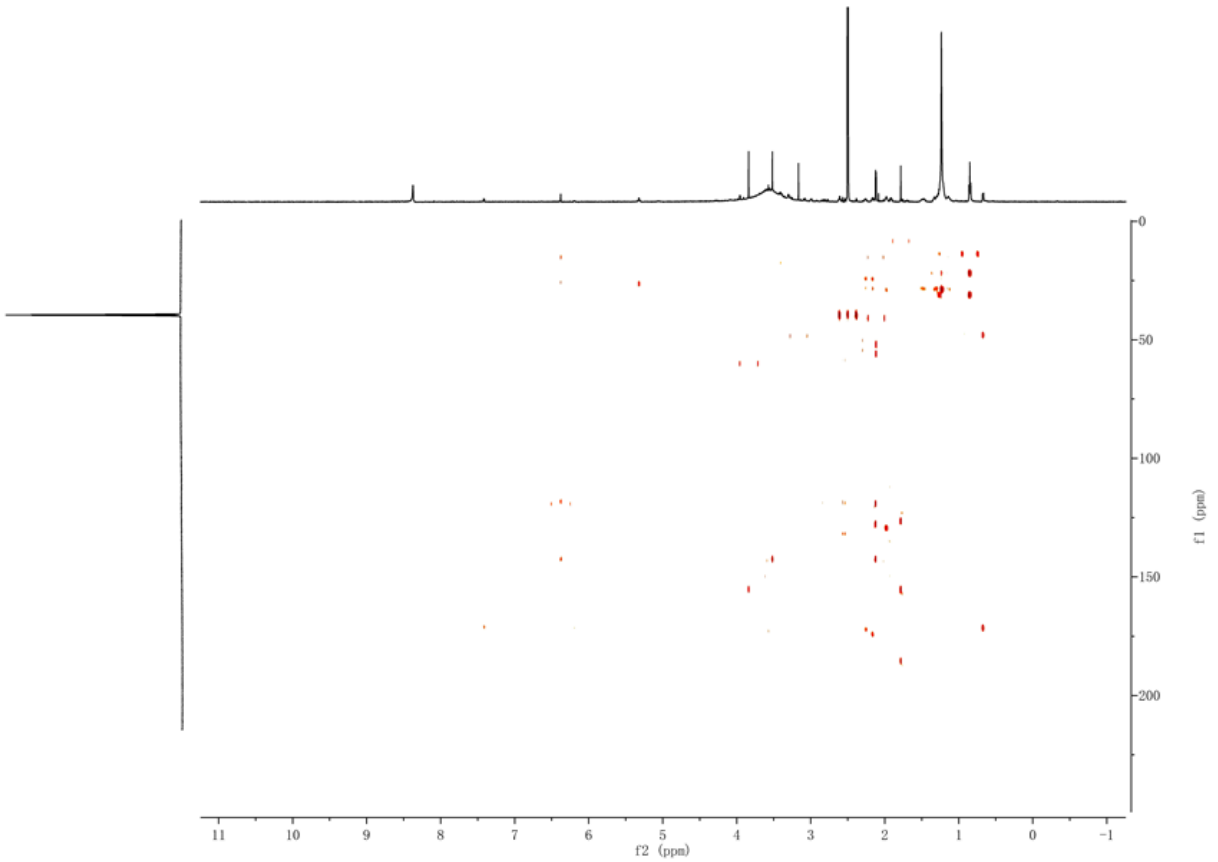


**Figure S12.** HMBC spectrum of compound **20** in DMSO-$d_6$.
